# Localized *in vivo* prodrug activation using radionuclides

**DOI:** 10.1101/2024.08.02.606075

**Authors:** J.M. Quintana, F. Jiang, M. Kang, V. Valladolid Onecha, A. Könik, L. Qin, V.E. Rodriguez, H. Hu, N. Borges, I. Khurana, L.I. Banla, M. Le Fur, P. Caravan, J. Schuemann, A. Bertolet, R. Weissleder, M.A. Miller, T.S.C. Ng

**Affiliations:** Center for Systems Biology, Massachusetts General Hospital Research Institute; Department of Radiation Oncology, Massachusetts General Hospital; Department of Imaging, Dana-Farber Cancer Institute; Office of Radiation Safety, Massachusetts General Hospital; Joint Program in Nuclear Medicine, Harvard Medical School; Department of Radiology, Massachusetts General Hospital, and Harvard Medical School; Institute for Innovation in Imaging, Massachusetts General Hospital; Department of Systems Biology, Harvard Medical School

**Keywords:** Theranostics, radioligand therapy, sustained drug delivery, combination chemoradiotherapy, FAPI

## Abstract

Radionuclides used for imaging and therapy can show high molecular specificity in the body with appropriate targeting ligands. We hypothesized that local energy delivered by molecularly targeted radionuclides could chemically activate prodrugs at disease sites while avoiding activation in off-target sites of toxicity. As proof-of-principle, we tested whether this strategy of “**RA**dionuclide **i**nduced **D**rug **E**ngagement for **R**elease” (**RAiDER**) could locally deliver combined radiation and chemotherapy to maximize tumor cytotoxicity while minimizing exposure to activated chemotherapy in off-target sites.

**Methods:** We screened the ability of radionuclides to chemically activate a model radiation-activated prodrug consisting of the microtubule destabilizing monomethyl auristatin E caged by a radiation-responsive phenyl azide (“caged-MMAE”) and interpreted experimental results using the radiobiology computational simulation suite TOPAS-nBio. RAiDER was evaluated in syngeneic mouse models of cancer using fibroblast activation protein inhibitor (FAPI) agents ^99m^Tc-FAPI-34 and ^177^Lu-FAPI-04, the prostate-specific membrane antigen (PSMA) agent ^177^Lu-PSMA-617, combined with caged-MMAE or caged-exatecan. Biodistribution in mice, combined with clinical dosimetry, estimated the relationship between radiopharmaceutical uptake in patients and anticipated concentrations of activated prodrug using RAiDER.

**Results:** RAiDER efficiency varied by 250-fold across radionuclides (^99m^Tc>^177^Lu>^64^Cu>^68^Ga>^223^Ra>^18^F), yielding up to 1.22µM prodrug activation per Gy of exposure from ^99m^Tc. Computational simulations implicated low-energy electron-mediated free radical formation as driving prodrug activation. Clinically relevant radionuclide concentrations chemically activated caged-MMAE restored its ability to destabilize microtubules and increased its cytotoxicity by up to 600-fold compared to non-irradiated prodrug. Mice treated with ^99m^Tc-FAPI-34 and caged-MMAE accumulated up to 3000× greater concentrations of activated MMAE in tumors compared to other tissues. RAiDER with ^99m^Tc-FAPI-34 or ^177^Lu-FAPI-04 delayed tumor growth, while monotherapies did not (*P*<0.03). Clinically-guided dosimetry suggests sufficient radiation doses can be delivered to activate therapeutically meaningful levels of prodrug.

**Conclusion:** This proof-of-concept study shows that RAiDER is compatible with multiple radionuclides commonly used in nuclear medicine and has the potential to improve the efficacy of radiopharmaceutical therapies to treat cancer safely. RAiDER thus shows promise as an effective strategy to treat disseminated malignancies and broadens the capability of radiopharmaceuticals to trigger diverse biological and therapeutic responses.

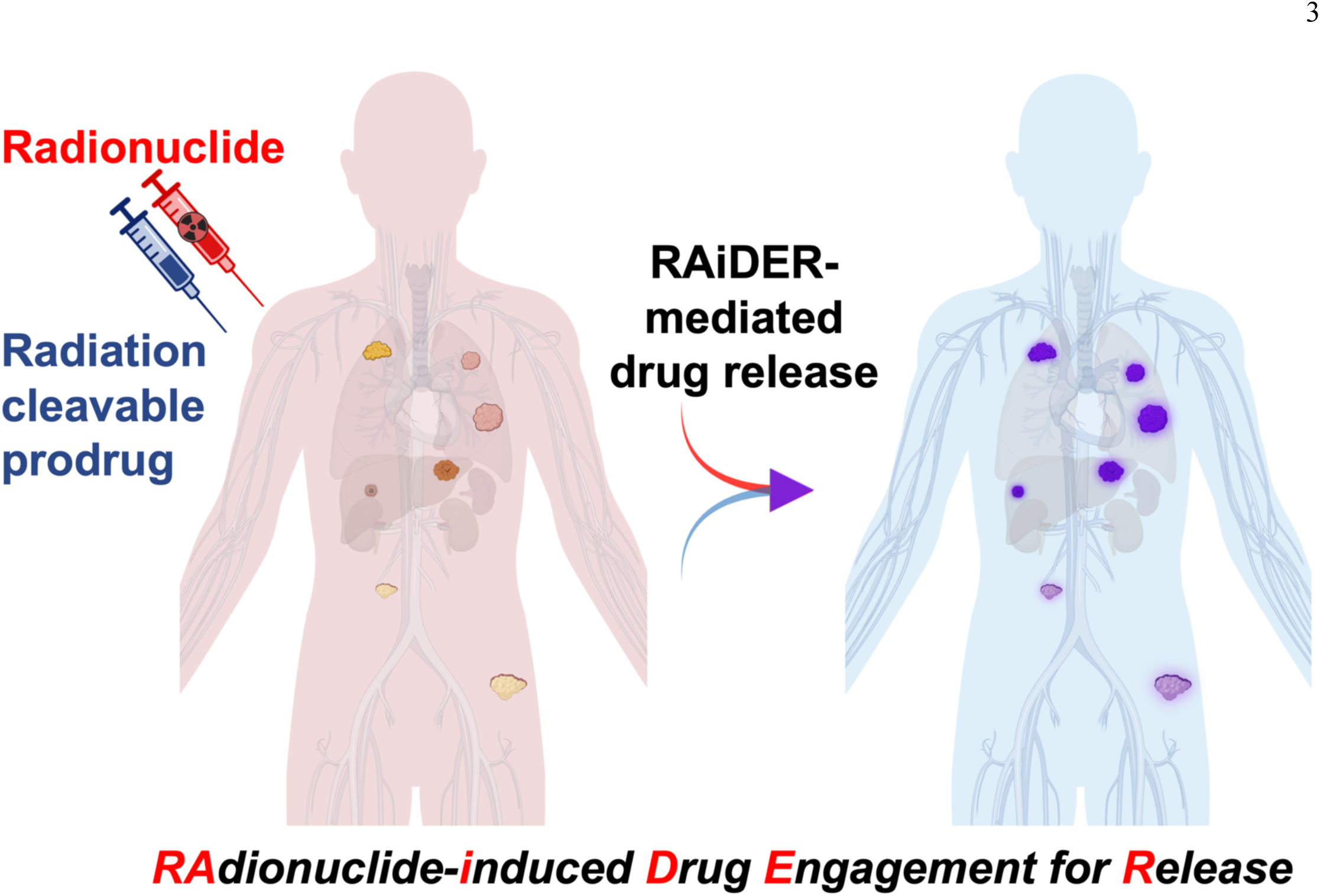

## Introduction

Radiopharmaceutical therapy (RPT) is becoming a critical pillar in cancer treatment, exemplified by FDA approvals of ^177^Lu-PSMA-617(*1*) and ^177^Lu-DOTATATE(*2*) for managing advanced-stage prostate and neuroendocrine tumors. Encouraging Phase III trials (NCT04689828, NCT03972488) are poised to support extending RPT indications to treatment settings earlier in the disease course(*3,4*). However, current RPT agents cannot mediate complete disease control in all patients. As with other therapy classes, RPT must be combined with other treatments for effective management of advanced disease(*5,6*).

Systemic chemo/immunotherapies are being actively tested with RPT (e.g. NCT03805594, NCT02358356, NCT05247905). As standalone therapies, they form the mainstay of treatment in patients with disseminated disease(*7*). While chemo/immunotherapies are often effective for tumor control, dose-limiting toxicities to off-target body sites constrain therapeutic indices(*8*). In some cases, over 50% of patients treated experience high-grade toxicities(*9*), exacerbated when multiple drugs are used in combination(*10*) or when other medical conditions exist. This results in dose reduction or treatment cessation, reducing treatment efficacy and potentially contributing to treatment resistance(*10*). Therefore, the ability to selectively deliver and activate potent drugs in tumors while minimizing their impact elsewhere in the body would be highly desirable.

Numerous drug delivery strategies have been developed to expand the range over which therapeutic cargoes can safely elicit effective responses(*11*). Examples include antibody-drug conjugates (ADCs) and nanomedicines designed to improve pharmacokinetics and payload delivery to tumors(*12,13*), prodrugs that release activated drugs in the presence of enriched enzymes or hypoxia(*14*), intraarterial and direct tumor delivery, and strategies using externally delivered triggers such as ultrasound to control drug activity(*15*). While such approaches can increase tumor target specificity and, in some cases, efficacy, unintended off-target activity remains a challenge. Inevitable accumulation in clearance tissues(*16*), heterogeneity of endogenous physiochemical properties within the tumor microenvironment, and premature drug activation due to chemical instability represent challenges for existing drug delivery approaches(*17,18*).

Ionizing radiation shows promise for triggering prodrug activation locally in tumors and addresses some of the above limitations(*19–22*). In principle, radiation-labile caging moieties enable precise spatiotemporal chemical activation of otherwise inactive prodrugs using external beam irradiation. A benefit of this approach over other prodrug strategies is that activation is bioorthogonal since drug release is triggered only in the presence of ionizing radiation. This approach can enhance release of activated chemotherapy within the tumor and block tumor growth without detectable systemic toxicities preclinically(*19,21,23*).

Translation of this approach is attractive for loco-regional or oligometastatic disease when using external beam radiation, but impractical for multifocal metastases(*24*). Appropriation of radionuclides, as used for RPT or other nuclear medicine examinations, for prodrug activation would extend the radiation drug activation paradigm for treating disseminated disease. Furthermore, when combined with RPT, local drug activation could, in principle, combine with radiotoxic and anti-tumor immunogenic effects of the radionuclide itself, a potentially powerful synergy(*3,5*). While examination of radiolysis in the context of maintaining radionuclide-conjugate stability for imaging and therapy has been explored(*25,26*), the application of a radionuclide-mediated drug activation approach remains limited(*27*).

This study assesses the feasibility of the “**RA**dionuclide **i**nduced **D**rug **E**ngagement for **R**elease” (RAiDER) approach(Fig. 1) for prodrug activation using a recently developed chemical strategy for long-circulating radiation-labile drug conjugates. Results show that radionuclides commonly used for diagnostic and therapeutic nuclear medicine can trigger intratumoral drug release from prodrugs without systemic toxicity nor off-target drug activation. Dosimetry guided by clinical nuclear medicine protocols suggests that the dose required for RAiDER is feasible in patients. This work thus offers a framework and proof-of-principle of how radionuclides can mediate localized prodrug activity in vivo.

**FIGURE 1.**
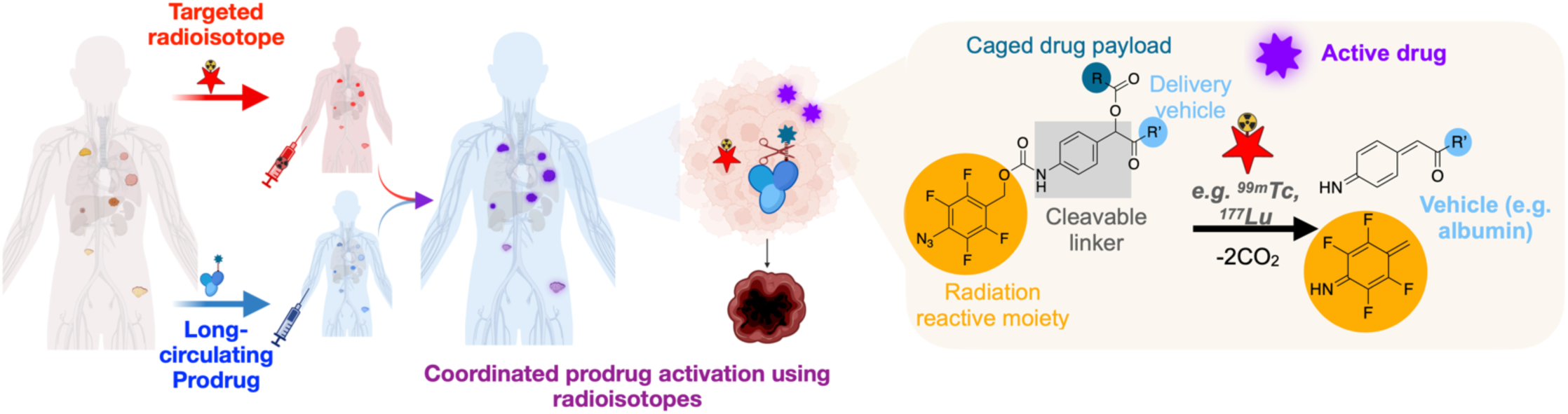
The RAiDER (“RAdionuclide induced Drug Engagement for Release”) concept. Targeted radionuclides accumulate in tumor tissues, where they locally deliver radiation to chemically activate caged prodrugs. Radionuclide- mediated prodrug activation occurs through reduction of a phenyl azide caging moiety(orange), leading to linker self-immolation and release of the active drug payload(purple) from its drug delivery vehicle(blue), which in this work is long-circulating serum albumin. Consequently, locally activated therapeutic payloads combine with ionizing radiation to maximize tumor cytotoxicity while sparing off-target tissues. Created with BioRender.com.

## Methods

Detailed experimental methods are presented in Supplemental Data. All animal research was performed under guidelines and approval from the local Institutional Animal Care and Use Committee.

## Results

### Synthesis and chemical characterization of long-circulating radiation-activated prodrugs

Since therapeutic radionuclides often exhibit long half-lives(e.g.^177^Lu t_1/2_:6.7 days) and deliver radiation over multiple weeks in patients, RAiDER was developed using a recent design for long-circulating prodrugs(*23*). This takes advantage of the long circulating half-life of serum albumin(human t_1/2_, ∼3 weeks), its preferential uptake across multiple tumor types, and consists of therapeutic payloads covalently conjugated to albumin via a radiation-labile para-azido-2,3,5,6- tetrafluorobenzyl(pATFB) moiety and self-immolating linker(*23,28*). Monomethyl auristatin E served as the model therapeutic payload using this strategy(Alb-caged-MMAE). A novel caged version of the topoisomerase inhibitor exatecan(Alb-caged-exatecan) was developed and used to assess the generalizability of the approach in some experiments(Schemes S1-3, Fig.S1-2). Constructs were stable in PBS(pH 7.4) for 4 weeks at 4°C(Fig.S3).

### Multiple radionuclide types can efficiently mediate RAiDER

The ability of commonly used radionuclides in clinical nuclear medicine to induce radiation-labile prodrug activation was evaluated in vitro. Alpha, electron/positron, Auger, and gamma emitters were assessed across preclinical and clinically relevant activities(Fig. 2A, Table S1). Results show roughly linear relationships between caged-MMAE activation and total dose imparted by each radionuclide. However, drug release efficiency between radionuclides was variable(Fig. 2B). ^99m^Tc and ^177^Lu showed highest efficiencies; ^99m^Tc was ∼7-fold more efficient than external beam radiation while ^177^Lu performed similarly. This was followed by other beta and alpha-emitting isotopes(^64^Cu, ^68^Ga, ^223^Ra). The near-pure positron-emitting isotope, ^18^F, was much less efficient. Diverse radionuclides thus elicit meaningful drug release using the pATFB radiation- labile caging moiety.

**FIGURE 2.**
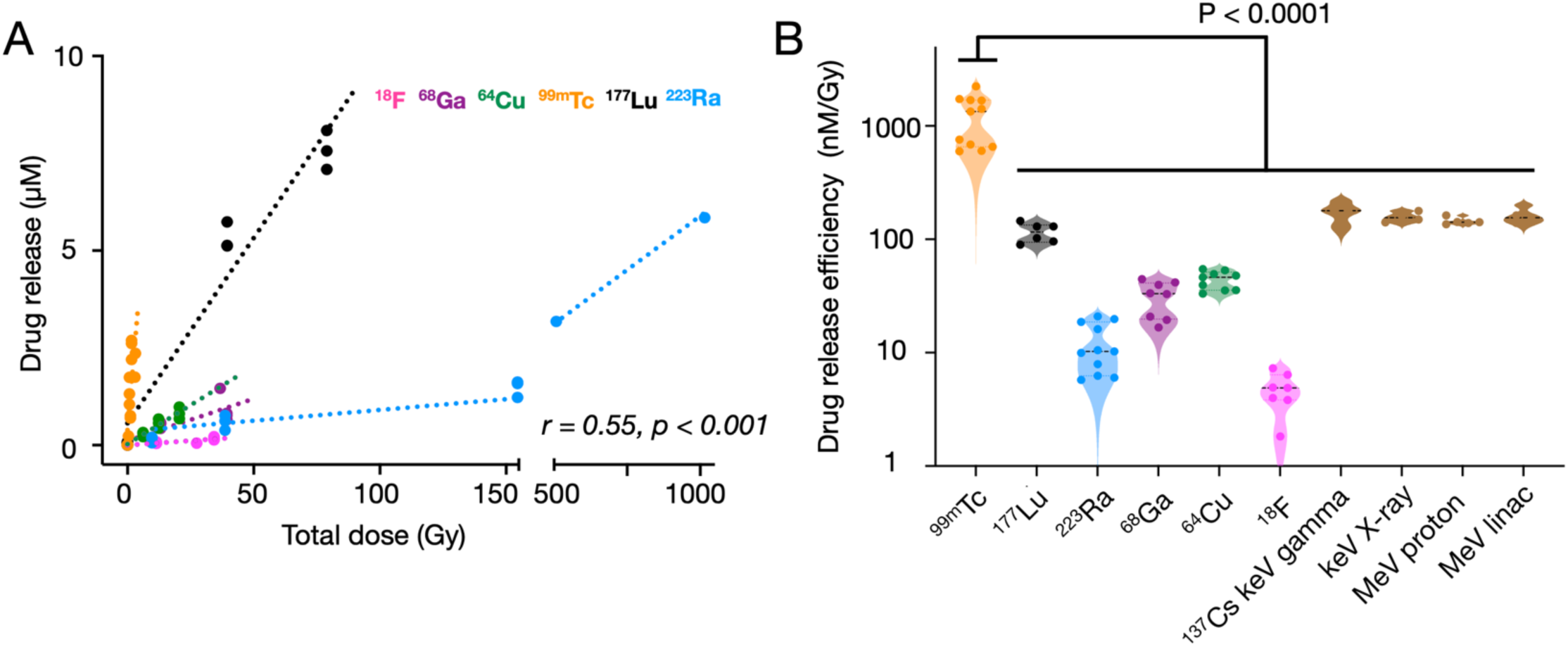
Radionuclide-mediated drug release from caged-MMAE. (A) 10µM caged-MMAE was exposed to different radionuclides at varying activities, with corresponding total doses released as estimated using TOPAS-nBIO(*34*); prodrug activation was monitored by LCMS/HPLC. (B) Drug release efficiencies across different radionuclides and external beam modalities are reported as nM-activated drug per Gy. The highest release efficiency(^99m^Tc) was compared to the other isotopes and modalities(mean±S.E., One-way ANOVA with Tukey’s multiple comparisons test, compared to ^99m^Tc). Please note that some values for ^137^Cs irradiation were previously presented(*23*) and reproduced here to facilitate comparison. Pearson correlation coefficient(r) was calculated across all data points.

### In vitro cytotoxicity of isotope-released prodrugs

Cytotoxicity of the isotope-released drug was tested in vitro in a panel of aggressive murine and human cancer models spanning across colon(MC38), prostate(LnCAP), anaplastic thyroid(TBP3743)(*29*), fibrosarcoma (HT1080), pancreatic(iKRAS)(*30*), and ovarian cancers(BBPNM)(*31*). Many of these diseases are less responsive to conventional chemotherapy(*29*). ^99m^Tc was used due to its high drug-activating efficiency. Alb-caged-MMAE activation by ^99m^Tc elicited up to a 600-fold increase in drug toxicity as measured by a 72h cell proliferation assay(Fig. 3A, S4, Table S2). ^99m^Tc alone was not cytotoxic at activities used for drug activation(Fig.S5). Clonogenic assays revealed that ^99m^Tc-activated Alb- caged-MMAE significantly blocked colony formation(Fig. 3B). Consistent with external irradiation(*23*), ^99m^Tc-activated Alb- caged-MMAE restored the ability of MMAE to destabilize cellular microtubules and induce apoptosis, as seen by immunofluorescence staining of alpha-tubulin and TUNEL, respectively(Fig. 3C). Therefore, radionuclide-activated caged- MMAE elicits biological effects in vitro consistent with the intended chemical prodrug release.

**FIGURE 3.**
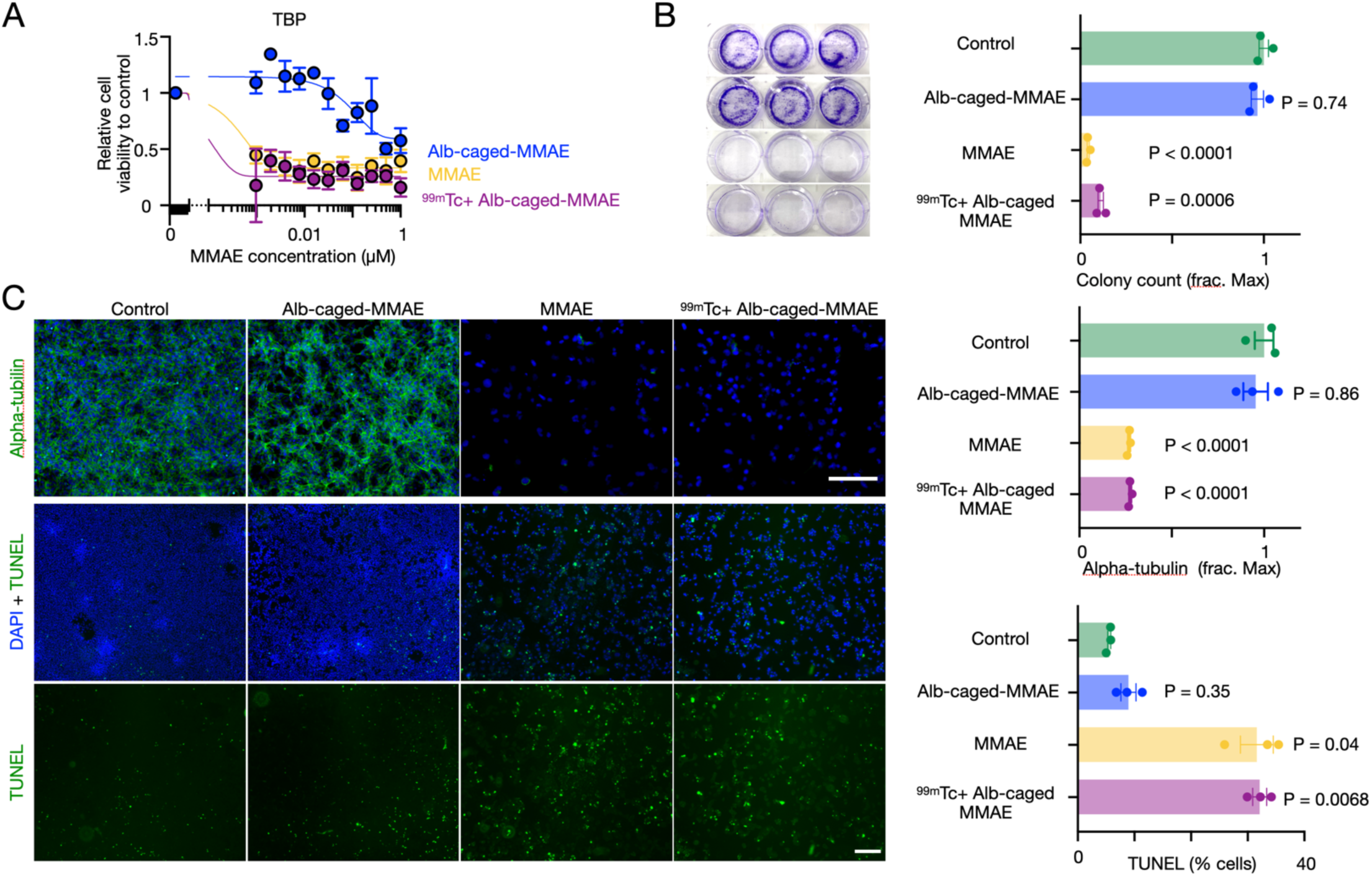
RAiDER restores biological prodrug activities in vitro. (A) Cytotoxicity of Alb-caged-MMAE, 99mTc, and free MMAE on TBP3743 anaplastic thyroid cancer, measured 72h post-treatment by a resazurin-based assay(n=3, mean±S.E.). (B) Representative images and quantification of TBP3743 colony formation(mean±S.E., 2-way-ANOVA with Geisser-Greenhouse correction). (C) Representative images of microtubule immunofluorescence(top) and apoptosis by TUNEL(bottom) 24h after treatment in TBP3743 cells(mean±S.E., 2-way-ANOVA with Geisser-Greenhouse correction, scale bars = 200µm).

### RAiDER depends on energy-specific free-radical formation

Free-radical products of water radiolysis including superoxide, hydroxyl radicals, and hydrated electrons underlie many radiation-dependent chemical reactions, including pATFB-uncaging and radiopharmaceutical stability(*25,26,32,33*). Therefore, we assessed whether radical scavengers could inhibit RAiDER, finding that gentisic acid and ascorbic acid could reduce caged-MMAE prodrug activation by ^99m^Tc(Fig.S6A). These data confirm RAiDER depends on availability of free radical species generated by ionizing radiation.

Free-radical generation from water radiolysis is effected by direct ionization from initial radionuclide decay events, such as from photoelectric effect or Compton scattering in the case of gamma/X-rays but more predominantly by indirect ionizations from secondary electron cascades with progressively lower energies along the ionization track(*34,35*). The spatial extent of these cascades depends on the initial energy of the electron from the original ionization. Radionuclides releasing high-energy positrons/electrons, such as ^18^F, would be expected to generate ionization events over a broader volume at lower concentrations than nuclides generating lower energy initial electrons(*36*). The higher concentration of ionization cascades for the latter may improve the probability of local reaction with macromolecules(*37*). These assertions(Fig. 4A) were tested using TOPAS-nBio, a Monte-Carlo simulation platform that examines radiobiology processes at sub-cellular scales(*34*). Simulations examining the number of electrons generated by radionuclides across different energy windows showed the best correlation with observed drug release across all nuclides for 100-110keV electrons(r=0.89, *P*<0.0001, Fig. 4B, S6B). As low-energy electrons (LEE, <30eV) are implicated in localized free-radical formation(*34,38*), we additionally simulated the dose deposited by LEE within nanoscopic volumes (∼10nm to reflect the hydrodynamic radius of Alb-caged-MMAE that would enable radicals to chemically interact with the prodrug), finding that the dose delivered by LEE at this scale correlated better with drug release than the total dose imparted across all radionuclides(Fig. 4C, 0.91 vs. 0.55, *P*<0.0001).

**FIGURE 4.**
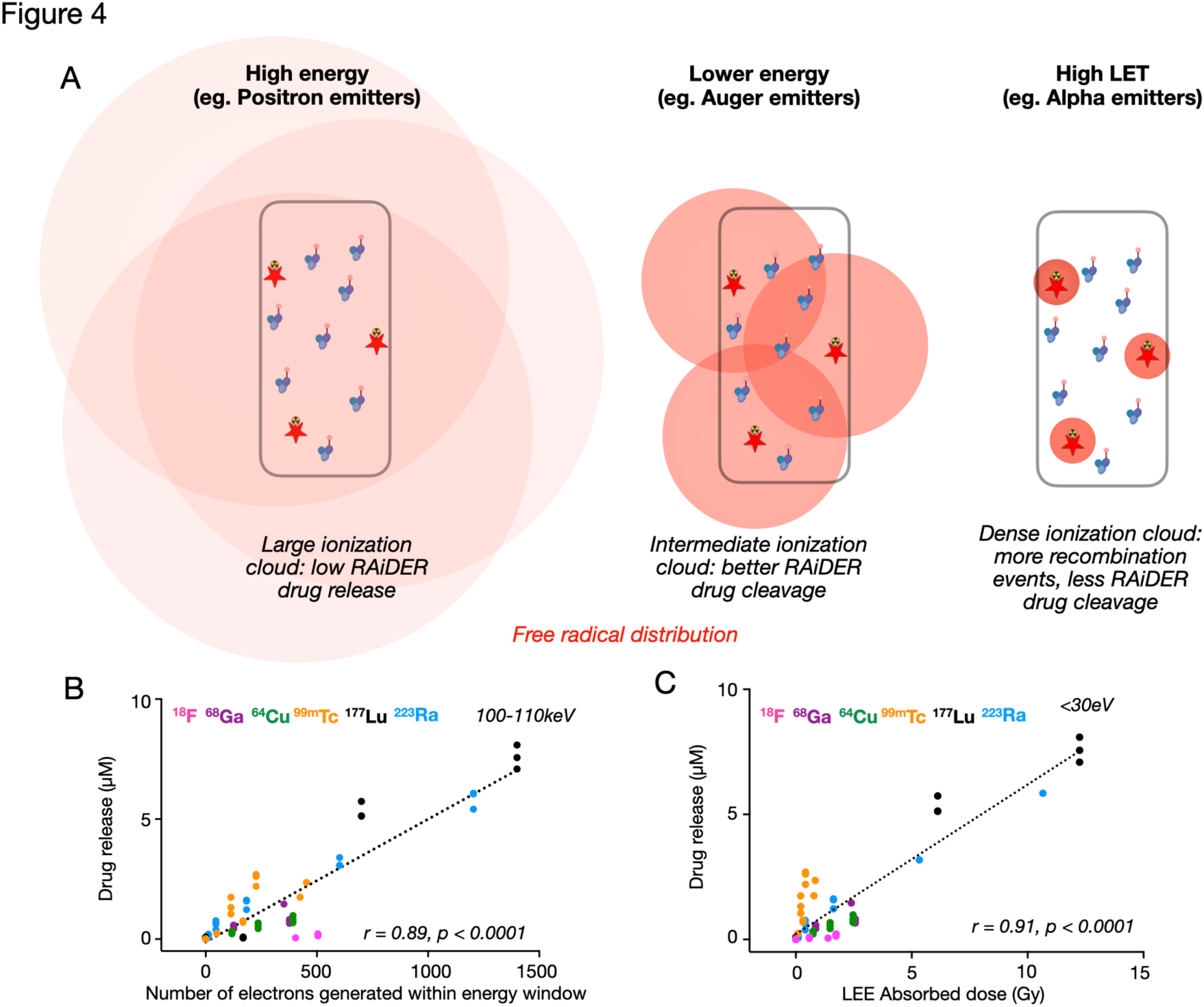
Computational modeling of radionuclide-mediated drug release using TOPAS-nBIO. (A) Conceptual mechanism to explain radionuclide-dependent RAiDER efficiencies. Red shading illustrates the spatial distribution and frequency of free-radical-generating ionization events from a given radionuclide within a vial. Ionization clouds interact with prodrug molecules most efficiently for isotopes emitting lower energy particles/photons (e.g. beta particles/Auger emitters, middle). (B) Correlation between observed drug release across radionuclides and their estimated number of electrons generated within the 100-110keV energy window. (C) Comparison of drug release efficiency with dose imparted by low energy electrons(LEE) across radionuclides. Pearson correlation coefficient(r) was calculated across all data points.

We experimentally tested the hypothesis that lower energy radiation enables more efficient drug release using external beam irradiation. X-rays from mammographic systems are less energetic(<50keV) than other conventionally used irradiation devices. We simulated energy spectra of two low-energy anode/filter combinations(W-Al/W-Rh) using SpekPy modelling X-rays from a mammographic system(*39,40*). Spectra generated from W-Al/W-Rh delivering ∼0.5Gy showed that both filters generate higher fluence at lower energies(≤60keV) than irradiation at 320keV(Fig.S7A, Table S3). This suggests that mammography-generated X-rays would lead to more efficient drug release, and this indeed was observed(Fig.S7B, *P*<0.0001). Together, results suggest that RAiDER is primarily driven by the local concentration of free-radicals from LEE and recombination effects of relatively low energy radiation.

### Radionuclide-mediated drug release in vivo

Next, we assessed the ability of RAiDER to mediate drug release in vivo, using the TBP3743 syngeneic mouse model of anaplastic thyroid cancer, an aggressive type of cancer characterized by poor clinical prognosis and traditional chemoradiotherapy resistance(*29*). Immunofluorescence indicated that TB3743 tumors express mouse fibroblast activation protein(muFAP, Fig.S8), and prior work suggests favorable tumor-to-background albumin uptake over 48h(*41*). Therefore, we treated mice bearing TBP3743 tumors with fluorescent Alb-caged-MMAE(^Cy5^Alb-caged-MMAE), ^99m^Tc- FAPI-34(*42,43*) administration 48h later, and tissue harvesting 24h later to assess biodistribution and prodrug activation(Figs.5, S9, S10). This showed highest accumulation and colocalization of both Cy5 fluorescence and ^99m^Tc- FAPI-34 in tumors. Other tissues examined showed uptake that was lower and/or discordant between the two agents. Selectivity for active MMAE accumulation was dramatically enhanced in tumors, 20-3000× higher than in other tissues(Fig.S11). Minimal caged-MMAE activation was seen without ^99m^Tc-FAPI-34(Fig. 5). Estimated dosimetry shows that in vivo prodrug activation by ^99m^Tc-FAPI-34 roughly matches values observed in vitro(Table S4(*44*)).

**FIGURE 5.**
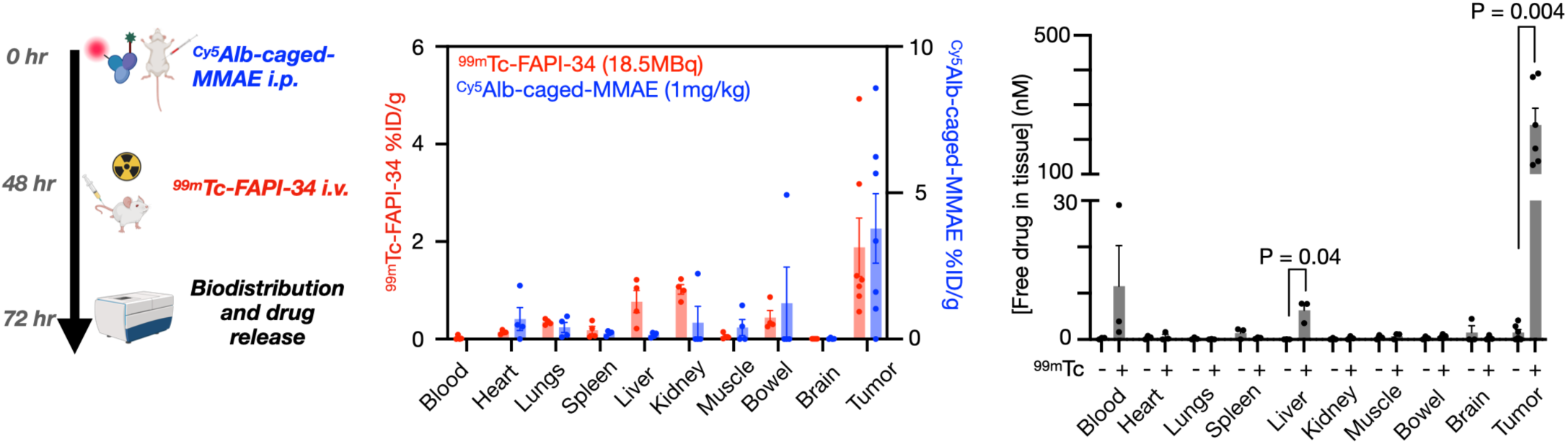
In vivo biodistribution and prodrug activation. Left: Experiment schematic of RAiDER biodistribution using B6129SF1/J mice bearing syngeneic TBP3743 tumors. Center: Biodistribution of fluorescent Alb-caged-MMAE, measured by tissue Cy5 fluorescence and 99mTc-FAPI-34 measured by gamma scintillation counting. Right: Biodistribution of activated Alb-caged-MMAE(with 99mTc-FAPI-34) compared to non-activated Alb-caged-MMAE as measured by LCMS/ HPLC. Data are means±S.E., n=3-4 mice, two-tailed t-test shown). Created with BioRender.com

To evaluate the generalizability of RAiDER, we performed analogous biodistribution assays using a syngeneic murine model of prostate-specific membrane antigen (PSMA)-expressing prostate cancer(RM1.PSMA, Fig.S12). As a proof-of-principle, we treated tumor-bearing mice with Alb-caged-exatecan and ^177^Lu-PSMA-617(Fig.S9(*45*)), which is used clinically to treat PSMA+ metastatic prostate cancer. ^177^Lu-PSMA-617 accumulated in tumors and triggered the local accumulation of activated exatecan more selectively in tumors compared to when free exatecan was administered. Taken together, biodistribution analysis indicates RAiDER efficiently and selectively releases active drugs at targeted tumor sites and is generalizable across therapeutic payloads, disease models, radionuclides, and their targeting strategies.

### RAiDER improves radionuclide efficacy to slow tumor progression

We next evaluated whether in vivo radionuclide-mediated prodrug activation could translate into detectable effects on disease progression. TBP3743 tumor-bearing mice were treated with Alb-caged-MMAE, ^99m^Tc-FAPI-34, or their combination, and tumor growth was monitored. Tumor growth was delayed with combination treatment compared to monotherapy/control treatments(Fig. 6A) and 2-way ANOVA indicated prodrug/radiation synergy(2-way ANOVA interaction *P*<0.006). No noticeable toxicity was observed, as evidenced by mouse weight changes during treatment, blood biomarkers, and histology(Fig.S13, S14). Of note, equimolar doses of free MMAE are known to be toxic in mice(*23*).

**FIGURE 6.**
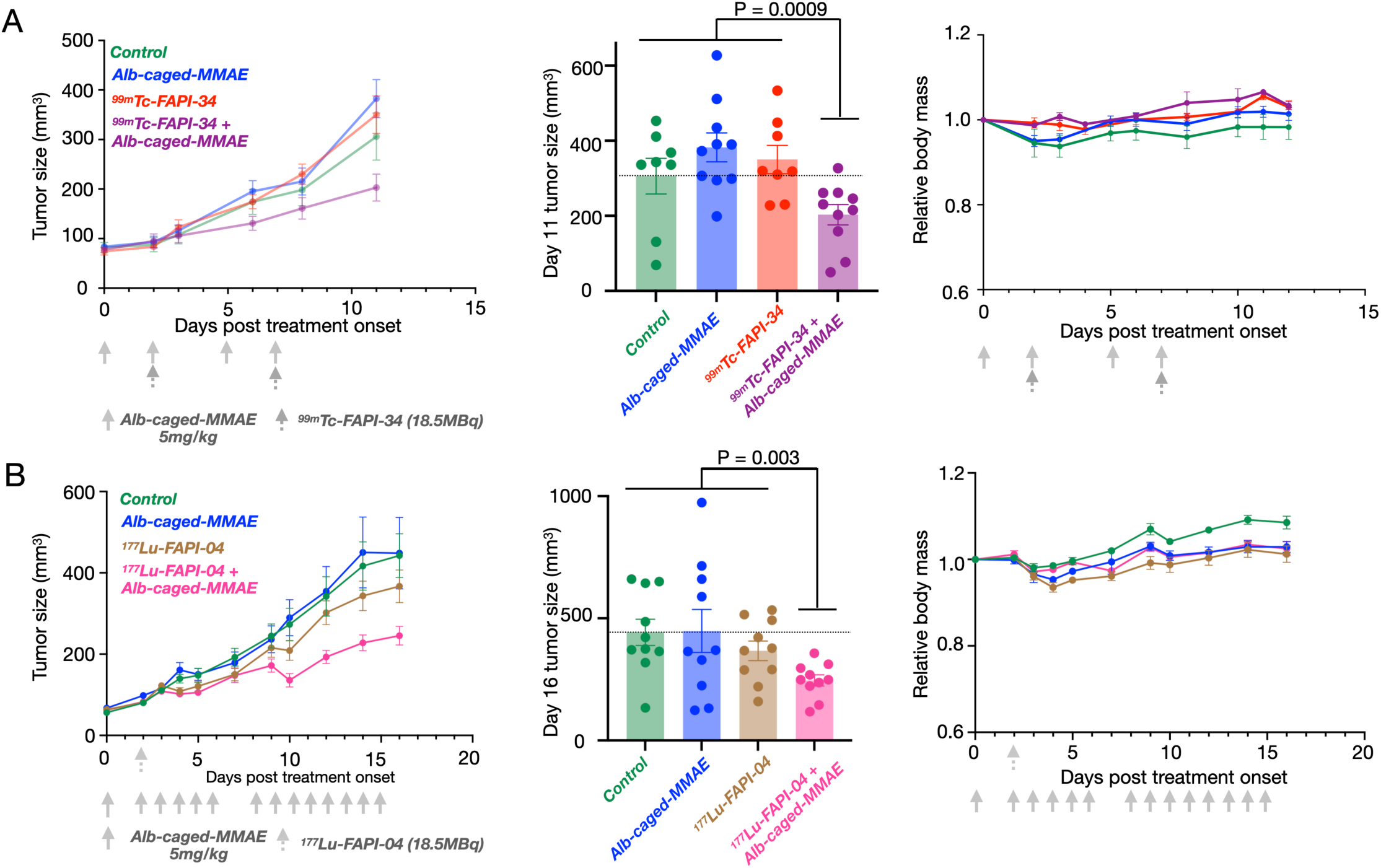
RAiDER enhances the ability of targeted radionuclides to block tumor growth. (A-B) Mice bearing TBP3743 tumors were treated with Alb-caged-MMAE(5mg/kg per dose) and either 99mTc-FAPI-34(18.5MBq per dose, A, n=18 total mice, 36 total tumors) or 177Lu-FAPI-34(18.5MBq, B, n=20 total mice, 40 total tumors). Caliper measurements of tumor growth are shown over time (left) and with individual tumor sizes on the indicated day (middle, two-tailed Mann-Whitney tests compare combination vs. pooled control/monotherapies). Mouse mass measured during treatment time course (right). Data displayed as mean±S.E.

We additionally tested RAiDER using ^177^Lu-FAPI-04(*46*), using a single injection to account for ^177^Lu’s long half-life. RPT- treatment significantly delayed tumor growth (2-way ANOVA term for RPT P<0.02). This was more pronounced when used in combination with Alb-caged-MMAE(Fig. 6B; compared to control/monotherapy P=0.003), without detectable toxicity. Together, these results demonstrate that RAiDER can safely improve radiopharmaceutical efficacy in a mouse model of aggressive cancer.

### Clinical dosimetry is compatible with RAiDER

Feasibility of RAiDER in patients was estimated using clinical dosimetry. Expected drug release in prostate and neuroendocrine tumors in patients treated with ^177^Lu-PSMA-617 and ^177^Lu-DOTATATE, respectively, was estimated based on published tumor dosimetry(*47,48*) and drug activation efficiencies shown in Fig. 2. Tumor accumulation of caged prodrugs was assumed not rate-limiting based on its observed biodistribution(Fig. 7) and pharmacokinetic modeling(*23*). Dosimetric analysis showed that radiopharmaceutical uptake in all lesions examined would be sufficient to mediate drug release well above the reported in vitro IC_50_ values of MMAE that block cancer cell proliferation(Figs.3, S4). Moreover, the standardized uptake values of these lesions are well within range of those seen in typical patients receiving radiopharmaceutical therapy.

**FIGURE 7.**
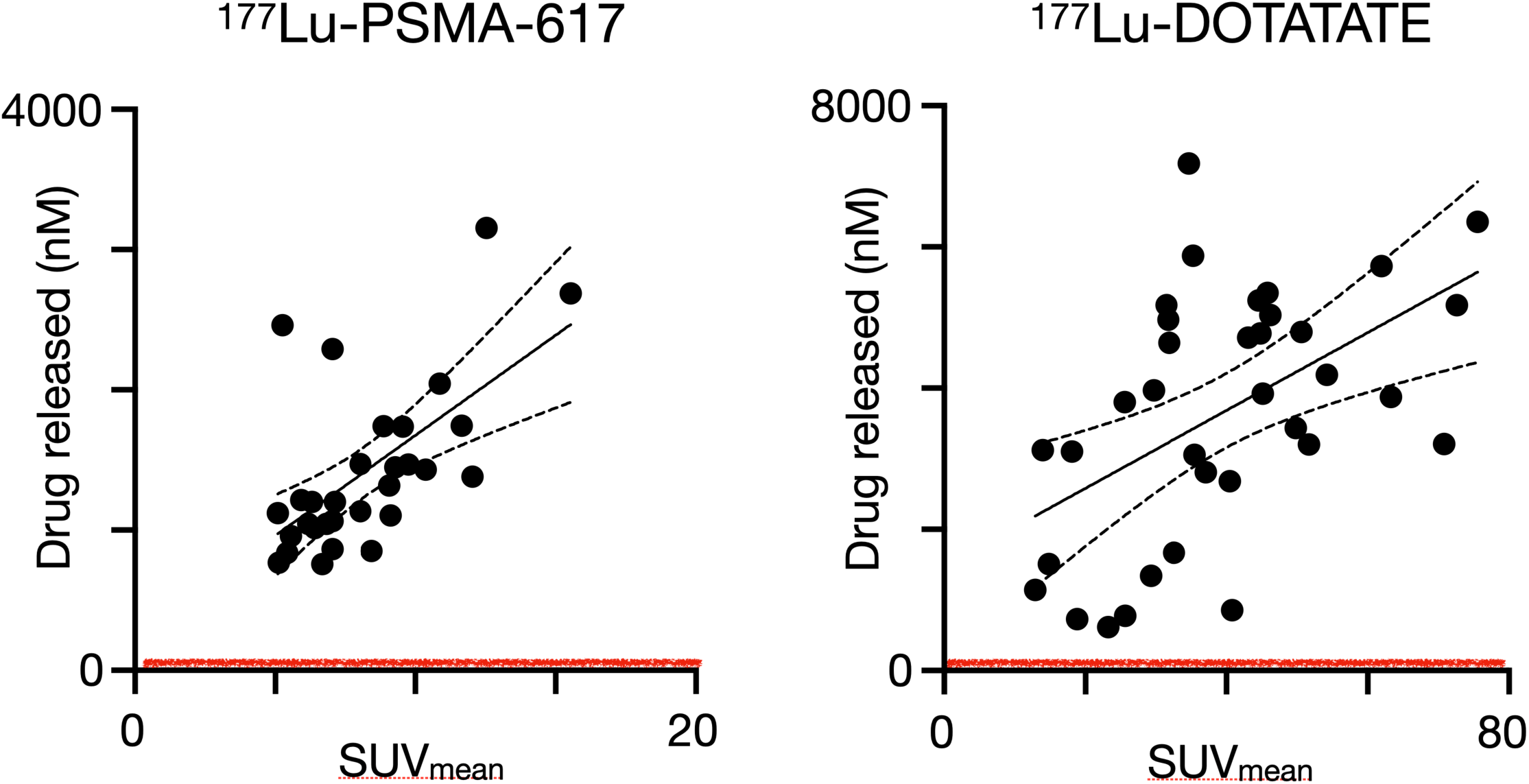
Estimating tumoral drug release in patients using clinical dosimetry. Lesion standardized uptake values (SUV) derived from 68Ga-PSMA-PET and 68Ga-DOTATATE-PET in prostate and neuroendocrine cancer patients receiving 177Lu-PSMA-617(47) and 177Lu-DOTATATE(48) respectively were compared to expected intratumoral active drug release(calculated based on release efficiencies in Fig. 2). Absorbed dose within each individual lesion was calculated in those previous studies using 177Lu SPECT/CT, demonstrating good correlation with PET-SUV. Red line denotes the IC50 estimated for aggressive cancers in this study (Table S2).

Extrapolation of these observations to other radiopharmaceuticals was assessed using MIRDcalc(*49*), using published pharmacokinetic data of agents currently in human use(*50–52*). As expected, the high linear energy transfer(LET) alpha-emitting isotopes would deliver significantly higher tumor dose per unit of activity compared to beta- and Auger-emitting isotopes regardless of the delivery vehicle(Fig.S15, Table S5, 2-way ANOVA with Tukey’s multiple comparison correction, *P*<0.02). Scaling these values to clinically used activities(alpha: 20MBq, beta: 7400 MBq, Auger: 1110 MBq), followed by an estimation of drug release based on the tumor dose delivered at these activities, shows that all classes of radionuclides, and especially ^177^Lu, can mediate drug release to enact meaningful(≥IC_50_) cytotoxicity across a range of tumor size and composition. This analysis suggests that RAiDER may be appropriate across a range of clinical applications, including for patients with borderline RPT lesion uptake that may be less responsive to RPT monotherapy.

## Discussion

This report presents RAiDER as a novel method to harness ionizing radiation imparted by radionuclides not only for their radiotoxic effects on tumor cells but also to mediate the localized release of complementary molecular therapy. Timed co-administration of caged drug payload(Alb-caged-MMAE) and its partner radiopharmaceutical enabled tumor colocalization of the two agents, enhanced tumor drug release, and reduced systemic exposure to free drug by 20-3000× in off-target tissues. Preclinical studies showed improved tumor control with RAiDER compared to RPT monotherapy without appreciable systemic toxicity.

Albumin-bound radiation-activated prodrugs were used in this proof-of-principle demonstration since serum albumin circulates for an extended period of time in the body via FcRn-mediated recycling. Its 66.5kDa molecular weight makes it smaller than antibodies and nanoparticles and promotes its ability to penetrate tissues, accumulating in cancer cells and phagocytes across multiple tumor types(*23,53*), facilitating translation of this approach across different malignancies. pATFB and alternative radiation-labile chemistries have been conjugated with other macromolecules or drug delivery vehicles, including nanoparticles or antibodies(*21,23*), likely applicable to RAiDER. Since the phenyl azide caging moiety is less sensitive to positron-emitting ^18^F, imaging versions of prodrugs could, in principle, be synthesized to provide a companion theranostic to guide treatment planning(*3*).

Prodrug activation using radionuclides has similarities and important differences compared to activation using external beam radiation. Cleavage efficiency by ^177^Lu or ^99m^Tc roughly matched or outperformed efficiencies from external beam radiation, respectively; radionuclides and external beam radiation activate drugs in a process dependent on the generation and availability of reactive radical species. Whereas external beam radiation is delivered at a high dose rate across discrete fractionations, radionuclides are generally retained at target lesions in the body and can facilitate more sustained prodrug activation(*54,55*). Different radionuclides showed different release efficiencies that were not directly related to LET, suggesting that not all ionized molecules translate into radicals available for prodrug activation. Further work remains to dissect specific mechanisms. Our in vitro quenching studies suggest that free radical-mediated linker cleavage mediates drug release, and simulation results suggest that a radionuclide’s ability to generate lower-energy electrons is more efficient for this process. We hypothesize that the efficacy of RAiDER may be a function of prolonged exposure to LEE-generating ionizing radiation within a localized volume. Ionized molecules can combine with free electrons, and radicals can recombine, depending on the local concentration of ionizations. These events may explain the reduced drug release yield for highly localized ionizing radiations, such as alpha emitters(*56*). Other factors, including tumor hypoxia, pH or the intracellular distribution of RPT and prodrugs, may impact activation efficiency and biological response. These interactions may also depend on the specific linker chemistry involved(*27*). Future studies characterizing the tumor microenvironment together with localized prodrug activation, controlled experiments with external beam radiation, and molecular simulations will further elucidate these mechanisms.

Heterogeneous tumor distribution is a barrier for antibodies, lipophilic drugs, and other targeted therapies. Incomplete drug distribution throughout the tumor can limit cytotoxicity and lead to treatment resistance(*57*). ADC linkers using enzymatic cleavage can be tuned to release their drug payload more efficiently across the tumor, including ADCs using MMAE(*58*). While this leads to better drug payload tumor penetration, this can result in unwanted drug release elsewhere, causing systemic toxicity. Moreover, enzymatic cleavage strategies, such as with cathepsins, require intracellular prodrug uptake to endolysosomes, which may not always occur efficiently. Radionuclide activation does not have this requirement, which may expand the types of molecular targeting strategies adopted for RAiDER, for instance, by targeting the extracellular matrix or surface-expressed tumor antigens, and is also better suited for extensive metastases than external beam radiation.

Implementation of RAiDER will depend on the pharmacologic properties of each component. This may be cancer-type dependent and require an in-depth understanding of the physiological response to a combination of RPT, and chemo/ immunotherapies. The optimal therapeutic index will likely be achieved with administration schemes that maximize tumor co-localization of RPT and prodrugs. For agents tagged with a short half-life nuclide such as ^99m^Tc, injection of a macromolecular-conjugated prodrug first allows sufficient drug accumulation before activation by the radionuclide. Conversely, longer half-life nuclides, such as ^177^Lu, can be injected before single or multiple prodrug doses to take advantage of prolonged tumor RPT retention(*54*). Choosing a radionuclide with better drug release efficiency may not always be advantageous if higher overall activity is injected using other nuclides for RPT. RAiDER optimization would benefit from a systems pharmacology approach combining experimental data and computation modeling. We have previously shown this approach to be helpful for other drug modalities(*59*).

Limitations of this study should be noted. We focused here on examining and maximizing tumor drug release using RAiDER. While results show relatively negligible drug release in off-target sites, a better understanding of the factors guiding radiation/prodrug interactions in normal tissues is warranted. Furthermore, MMAE and exatecan may not be the optimal drug combination for RAiDER. A broad palette of drugs can be used with radiation-labile linkers; identifying those that synergize with RPT is essential. These include immunotherapies and therapies targeting pro-tumorigenic signaling pathways activated by ionizing radiation(*60*). Systematic experimental assessment of other radionuclides(e.g., ^90^Y, ^131^I, ^225^Ac) in vitro and in vivo is also needed to augment our understanding of the integrated physical, chemical, and biological parameters that govern the efficacy of RAiDER.

In summary, we demonstrate that multiple clinically relevant radionuclides can control the spatial-temporal release of albumin-linked radiation-cleavable prodrugs in vitro and in vivo. Judicious choice of radiopharmaceutical and caged- prodrug combinations enable an effective combination systemic therapy approach with excellent therapeutic indices for disseminated disease. This work sets the stage for preclinical and clinical RAiDER evaluation across various malignancies.

## Acknowledgments

We thank Drs. Gregory Wojtkiewicz, David Lee, Kyle Stewart, Hamid Sabet, Allegra di Pietro, Michael Zurawski, Umar Mahmood, Pedram Heidari, Ciprian Catana, Pedro Brugarolas, and Xin Gao for helpful discussions and support.

## KEY POINTS

### QUESTION

Can radionuclides induce localized prodrug activation in vivo?

### PERTINENT FINDINGS

The ability of radionuclides to mediate the chemical activation of prodrugs was demonstrated across clinically relevant dosings of radionuclides. Molecularly targeted radionuclides could induce active drug release from long-circulating prodrug-conjugates preferentially in tumors over off-target sites in the body. Combination radiopharmaceutical therapy with a model prodrug (caged-MMAE) delayed tumor growth in an aggressive mouse model of anaplastic thyroid cancer without appreciable toxicity, whereas monotherapy did not.

### IMPLICATIONS FOR PATIENT CARE

The “**RA**dionuclide **i**nduced **D**rug **E**ngagement for **R**elease” (**RAiDER**) strategy is feasible in vivo, paving the way for designing effective combination radiopharmaceutical treatments with high therapeutic indices using this approach.

