## Supplemental Data for "Localized *in vivo* prodrug activation using radionuclides"

### Materials and Methods

***Synthesis of the radio-cleavable prodrug linker for RAiDER.***

Unless stated otherwise, all materials were used as received from commercial sources. N,N-diisopropylethylamine, N,N-dimethylformamide, bis(4-nitrophenyl) carbonate, lithium hydroxide, and HBTU were purchased from Sigma Aldrich (St. Louis, MO, USA). Hydrochloric acid (HCl), methanol, dichloromethane, and acetonitrile were purchased from VWR International (Radnor, PA, USA), while Monomethyl auristatin E (MMAE) and exatecan were purchased from MedChem Express (Monmouth Junction, NJ, USA). Maleimide-PEG4-amine trifluoroacetic acid salt was purchased from BroadPharm (San Diego, CA, USA) and $CDCl_3$ was purchased from Cambridge Isotope Laboratories (Tewksbury, MA, USA). Reaction mixtures were purified using a Biotage Sfar Bio C18 (300 Å, 10 g) on a BUCHI C-850 FlashPrep with a gradient composed of water (0.1% formic acid) and acetonitrile (0.1% formic acid) for reversed-phase chromatography. $^{1}H$, and $^{13}C$ NMR spectra were recorded on a Bruker AC-400 MHz spectrometer. High-performance liquid chromatography-mass spectrometry (HPLC-MS, LCMS) analysis was performed on a Waters instrument equipped with a Waters 2424 ELS Detector, a Waters 2998 UV-Vis Diode array Detector, a Waters 2475 Multi-wavelength Fluorescence Detector, and a Waters 3100 Mass Detector. Separations employed an HPLC-grade water/acetonitrile solvent gradient. Columns: XTerra MS C18 Column, 125, 5 µm, 4.6 mm X 50 mm column. All samples were run on an XTerra MS C18 column using a gradient of 5 to 95% acetonitrile in water (0.1% formic acid) over 1.5 minutes, followed by 95% acetonitrile for 0.5 minutes at a flow rate of 5 mL/min.

Caged-MMAE and caged-exatecan were synthesized as outlined in Scheme S1, guided by previous works(*1*). Briefly, the drugs were conjugated to the pATFB-SIL (para-azido-2,3,5,6-tetrafluorobenzyl self-immolative linker) via a carbamate linkage under basic conditions, followed by ester hydrolysis and amide coupling to install a maleimide anchor. The resulting prodrugs were characterized via LCMS and NMR. Synthetic details can be found below in the following section (Schemes S1-S3, Fig. S1, S2). Maleimide was reacted with the free cysteine on mouse serum albumin to yield albumin-conjugated caged prodrugs(*1*). The reactive fluorophore Cyanine5-NHS ester (Lumiprobe) was conjugated to serum albumin prodrug via amine coupling (DOL: 3.2) for biodistribution experiments (*1*). Conjugate stability was determined by treatment of the conjugates with tris(2-carboxyethyl)phosphine (TCEP, 10eq), which reduces the azide linker via a Staudinger reduction, leading to release of the free drug. Prodrug aliquots were purified at various time intervals via spin filtration through a 10kDa molecular-weight cutoff centrifugal filter (Amicon) 10,000 rcf for 7 min), and the free drug was quantified via LCMS to determine the presence of intact prodrug over 30 days.

***Prodrug activation using radionuclides in vitro*.**

Radionuclides examined in this study, their formulation and source, and the conditions used for their testing are outlined in Table S1. For in vitro studies, 10µM of caged-MMAE was incubated with varying activities of each isotope for defined periods (Table S1) in 2mL HPLC glass vials (Thermofisher, USA) and buffered with 10× PBS to achieve a volume of 1mL at pH7.4 and a final concentration of 1×PBS. Samples were kept at least 0.5cm apart during incubation at room temperature. No appreciable cross-vial mediated drug release was found in the control vials placed in this configuration in pilot studies. Samples for which the incubation time was shorter than the physical half-life of the radionuclide were frozen and stored in a liquid nitrogen tank until further analysis could be performed. This cryopreservation prevented further drug release from the pATFB linker (Fig. S3D). Chemical quenching studies were performed with gentisic and ascorbic acids (Millipore Sigma).

***Prodrug activation using external radiation.***

Prodrug activation was additionally tested with various methods of external beam radiation. Samples were prepared as above, except in 96 well plates diluted to a volume of 100µL. A total of five irradiation modalities were used:

*X-ray irradiation*: X-ray irradiation used an X-RAD320 cell and small animal irradiator with a 320keV energy and dose rate of 325±10cGymin$^{-1}$ at room temperature (Precision X-ray). Parameters for X-ray irradiation were 320kV, 12.5mA with 3mm Be (internal) and 2mm Al (external) filters applied.

*Gamma irradiation*: Irradiation was performed on a dual source $^{137}$Cs Gammacell 40 Exactor (Best Theratronics) with a dose rate of roughly 50cGymin$^{-1}$.

*Protons*: Proton beam irradiations were performed at the Francis H. Burr Proton Therapy Center with a single field impinging vertically on the samples. The samples were placed at the center of the spread-out Bragg peak (SOBP) with a range of 13cm and a modulation width of 7cm, resulting in a flat dose distribution at and around the sample with a linear energy transfer (LET) of ~2.3keVµm$^{-1}$ and a dose rate of ~0.5Gys$^{-1}$. The LET was estimated using Monte Carlo simulations with TOPAS, which was previously well-tested for this beamline(*2*).

*MV Linac*: Clinical energy (6MV) photon irradiations were performed at the MGH Clark Center using Varian Truebeam Linacs. A solid water phantom size of 5 or 10cm depth was used to simulate the internal scatter expected with biological tissues.

*Mammography*: This was performed with a Hologic 3Dimensions mammography system. Exposure dose was measured using a portable calibrated dosimeter (Raysafe X2) connected to a solid-state mammographic sensor (X2 MAM). If necessary, samples were stored at 4ºC until analysis. The linearity of drug release using this approach across different doses and multi-day experimental sessions was confirmed before additional experiments (Fig. S7C).

***Estimation of drug release.***

Samples (for tissues, an extraction procedure was performed prior to analysis as described below) were analyzed using LCMS (Waters instrument equipped with a Waters 2424 ELS Detector, Waters 2998 UV-Vis Diode array Detector, and a Waters 3100 Mass Detector) if the incubation time exceeded 10 half-lives of the relevant isotope or using an Agilent 1200 Series HPLC, with a multichannel-wavelength UV/Vis detector (G1365D), fluorescence detector (G1321A), and a flow-through γ-detector (γ-Ram, LabLogic, with a 35900E interface), using PBS mobile phase at a flow rate of 0.7 mL $min^{-1}$) if the sample was radioactive at the time of analysis. On both instruments, an Xterra MS C18 Column (Waters; 124A, 5 μm, 4.6 x 50 mm) column was used, with a gradient of 5-95% acetonitrile in water with 0.1% formic acid as the mobile phase. Drug concentration from LCMS and LC/UV samples were quantified by comparing the area under the curve from the ELSD, isolated mass chromatographs (+ESI 718.8 Da for MMAE, 436.4 Da for exatecan), or UV absorbance (210 nm for MMAE, 320 nm for exatecan) of each sample to a 7-point standard calibration curve (10 μM to 39 nM, Limit of Detection: ~15 nM). LCMS and UV standards were cross calibrated to allow for direct comparison.

***Cell culture***

Mouse cancer cell lines underwent mouse pathogen testing (IDEXX) before use, and all cells were routinely tested for mycoplasma contamination (Mycoplasma PCR test Kit, Applied Biological Materials). TBP3743 (murine anaplastic thyroid cancer(*3*)), HT1080 (human fibrosarcoma with and without human histone H2B-green fluorescent protein expression to visualize the nucleus (HT1080-H2B-GFP) (*1*)), BPPNM (murine ovarian cancer), LnCAP (human prostate cancer, ATCC), RM1.PSMA (murine prostate cancer with prostate-specific membrane antigen, PSMA, constitutively expressed using a PiggyBac transposase cassette, plasmid procured from VectorBuilder with original cell line obtained from ATCC), MC38 (murine colorectal cancer), and iKras (murine pancreatic cancer) cells were prepared as previously described(*4*). TBP3743, HT1080, RM1.PSMA, and MC38 were cultured in DMEM, while LnCAP used RPMI, all supplemented with 10%

fetal bovine serum (FBS) and penicillin/streptomycin (P/S) under an atmosphere of 5% $CO_2$. AC37 mouse cancer cells with doxycycline-inducible KrasG12D expression (iKras, a gift from Dr. Haoqiang Ying, MD Anderson Cancer Center, by way of Dr. Nabeel Bardeesy, MGH), derived from triple transgenic p48-Cre; ROSA26-LSL-rtTa-IRES-GFP;TetO-LSL-KrasG12D genetically engineered mouse model of pancreatic ductal carcinoma(*1*), were maintained in DMEM/Nutrient Mixture F-12 media (DMEM/F12, Invitrogen) supplemented with 2 μg/mL doxycycline (Sigma). The BPPNM cells(*5*) were cultured in DMEM supplemented with 1% insulin–transferrin–selenium (Thermo Fisher Scientific), epidermal growth factor (2 ng/mL), 4% heat-inactivated FBS (Thermo Fisher Scientific), and 1% P/S.

***Cytotoxicity assay of cleaved drugs.***

Cytotoxicity experiments were performed by seeding 3000 cells per well overnight in a 96-well plate (Corning) before adding each drug/conjugate. MMAE or prodrug (with or without exposure to $[^{99m}Tc]TcO_4^-$ for at least 60h to promote drug activation) were prepared at varying concentrations ([MMAE], 1μM and subsequent 2-fold dilution to 1nM) in media. Empty wells with only media or vehicle treatment served as controls. After adding the corresponding drug condition and a 72h incubation, PrestoBlue (ThermoFisher, USA) determined the number of live cells according to the provider's protocols.

***Radiotoxicity assay.***

3000 cells per well were seeded overnight in a 96-well plate (Corning) before adding $[^{99m}Tc]TcO_4^-$ at varying concentrations. Empty wells with only media or vehicle treatment served as controls. After a 72h incubation, PrestoBlue (ThermoFisher, USA) determined the number of live cells.

***Colony formation assay***.

$1 \times 10^4$ cell lines were placed in each well of 96-well plates and incubated overnight. The cells were treated with either a control solution or $^{99m}$Tc-activated caged-MMAE.  After 48h, the cells were harvested and reseeded into individual wells of a 6-well culture plate in fresh medium at 200 cells/mL and allowed to form colonies for 1 to 3 weeks. Once colonies had developed, they were fixed with 100% methanol for 20 minutes, stained with a crystal violet staining solution (Sigma) for 20 minutes at room temperature, and then rinsed with distilled water. Plating efficiency (PE) and survival fraction were calculated as previously described: Survival fraction = (number of colonies formed after treatment) / (number of cells initially seeded × PE) × 100%, plating efficiency (PE) = (number of colonies formed for untreated cells) / (number of cells initially seeded) × 100%. The plating efficiency for non-irradiated cells was determined to be 95%.

***Immunofluorescence and TUNEL assays***

1 × $10^4$ TBP3743 cells were seeded in each well of 96-well plates overnight. TBP3743 cells were then treated with vehicle, Alb-caged-MMAE (with and without $^{99m}$Tc-mediated cleavage), or MMAE at a concentration of 100nM. After 6h, the cells were washed three times with PBS, replaced with fresh cell culture media, irradiated with 10Gy X-rays, and incubated for 24h (for TUNEL marker staining) or 48h (for α-tubulin immunofluorescence). Cells were then fixed with 4% paraformaldehyde (PFA) for 30 minutes at room temperature (RT), washed in PBS, permeabilized in 0.5% Triton X-100 in PBS for 30 minutes, and blocked with a 10% normal serum for 20 mins at room temperature. α-tubulin Alexa Fluor 488 mouse monoclonal antibody (DM1A, 1:100, Invitrogen) was used for staining. Apoptotic cells were determined using a DeadEnd Fluorometric TUNEL System (Promega) according to the manufacturer’s instructions.

Fibroblast activating protein (FAP) expression in TBP3743 tumors was confirmed using immunofluorescence. Tumor samples were harvested and stored in 4% paraformaldehyde overnight and subsequently in sucrose 15% solution for 12h, then in sucrose 30% solution overnight, and finally stored in 1xPBS until sectioning. Intact tissue was prepared for sectioning by embedding using Optimal Cutting Temperature (OCT, Sakura) in a cryomold and frozen on dry ice. 10 μm cryosections were prepared on glass slides. Tissue sections were first blocked with blocking buffer (5% normal goat serum, 0.2% Triton X-100, 5% BSA 1× PBS) for 2h at RT. Samples were then incubated with a primary antibody (mouse fibroblast activation protein mFAP antibody, R&D system, monoclonal Rat IgG1 stock: 0.5 mg/ml, 1:50 dilution in wash buffer: 10% normal serum, 0.2% Triton x-100 0.2% in PBS) overnight at 4°C in a humidified staining chamber. Samples incubated with only the secondary antibody served as negative controls. Slides were then washed with PBS 3 times, followed by staining with a secondary antibody for 1h at RT (AF488 anti-rat IgG2b antibody, Biolegend, Mouse IgG1: 0.5 mg/ml. Clone: MRG2b-85, 1:200 dilution in wash buffer).

PSMA expression in RM1.PSMA cells was evaluated by immunofluorescence staining. 5 × $10^3$ RM1.PSMA cells were plated in a 96-well plate overnight before being fixed with 4% PFA. Cells were then blocked with blocking buffer at RT for 1.5h, followed by primary PSMA antibody incubation for 1h (anti-FOLH1, monoclonal Rabbit stock IgG1, Abclonal, #A9547, 1:200). Cells were washed 3 times with PBS, and then stained with a secondary antibody the solution (AF647 goat anti-rabbit IgG1, Abclonal, #AS075 1:200). HT1080-H2B-GFP cells stained similarly served as negative control.

All samples were mounted with VECTASHIELD® Antifade Mounting medium with DAPI to stain the nuclei and stored until imaging. Slide-mounted samples were imaged using a fluorescent microscope (Revolve, Discover Echo). Images were analyzed using ImageJ and/or CellProfiler. α-tubulin and TUNEL stains were quantified by calculating the positive signal normalized to the vehicle control.

***Preparation of $^{99m}$Tc and $^{177}$Lu labeled FAPI and PSMA targeting agents***

$^{99m}$Tc-FAPI-34, $^{177}$Lu-FAPI-04, and $^{177}$Lu-PSMA-617 used in this study were prepared as guided by prior reports (*6,7*). Precursors were commercially obtained (MedChemExpress). Reduction of $[^{99m}TcO_4]^-$ to $[^{99m}Tc(CO)_3]^+$ was performed using a formulation similar to the IsoLink kit(*8*), consisting of 2.85mg sodium tetraborate $Na_2B_4O_7$, 4.5mg sodium boranocarbonate $Na_2H_3BCO_2$, 7.15mg sodium carbonate $Na_2CO_3$ and 8.5mg sodium tartrate $Na_2C_4H_4O_6$, dissolved in 1mL ultrapure milliQ water (Millipore, Billerica, MA, USA) in a 10mL glass vial and degassed with nitrogen for at least 15 minutes. This was subdivided into aliquots of 260μL in microcentrifuge tubes with rubber ring-sealed screw caps (Nalgene, Rochester, NY, USA) under an anaerobic environment for further use. For labeling, up to 1GBq $[^{99m}TcO_4]^-$, generator elulate (100μL) was added to each tube and heated for 30min at 100°C. The microcentrifuge tube was then allowed to cool to RT, and the solution neutralized with 25μL of 1M HCl to pH approximately 7.5, giving a total volume of 125μL of $[^{99m}Tc(CO)_3]^+$.

The radiochemical purity (RCP) of the $[^{99m}Tc(CO)_3]^+$ produced was checked with thin layer chromatography (TLC) plates (3cm × 7.5cm, Merck, Darmstadt, Germany) using a mobile phase of 1% HCl in methanol. Plates were analyzed with a gamma ray radioTLC scanner (AR2000). $[^{99m}Tc(CO)_3]^+$ was then added to a mixture of 5μL of the individual FAPI-34 precursor (1mM in water). The reaction was then heated to 95°C for 20min.

Labeling completeness of all three radiopharmaceuticals was monitored by radioTLC (Fig. S9) and processed by solid-phase extraction, evaporation, and formulation with 0.9% saline before therapy or biodistribution experiments. iTLC was performed by loading each chelate solution (1μL) onto glass microfiber chromatography paper impregnated with silica gel (iTLC-SG, Agilent), then run in a solution of 0.2M sodium acetate (pH 6). The iTLC was then analyzed on an AR-2000 (Eckert & Ziegler) TLC scanner. The RCP of all agents were >95%.

***Mouse experiments.***

All animal research was performed under guidelines and approval from the local Institutional Animal Care and Use Committee. Mice were housed in a pathogen-free vivarium with controlled temperature, humidity, and light/dark cycling. B6129SF1/J (TBP) and C57BL6/J (RM1.PSMA) mice were purchased from the Jackson Laboratory (JAX).

***Biodistribution of caged prodrugs and co-administered radionuclide***.

*TBP3743 anaplastic thyroid cancer model*: 5-12-week-old female B6129SF1/J mice were inoculated with $5 \times 10^5$ TBP3743 cells subcutaneously. Once tumor reached ~100mm$^3$ roughly 10 days later, fluorescent Alb-caged-MMAE ($^{Cy5}$MSA-pATFB-MMAE) was injected intraperitoneally (1.4µmol/kg; 1mg/kg free MMAE equivalent). A subset of these mice was injected with 18.5MBq $^{99m}$Tc-FAPI-34 intravenously via retro-orbital injection 48h later (n = 3-4 per treatment condition). Tissues were harvested from the mice after an additional 24h. Tissues from additional mice obtained 8h after $^{99m}$Tc-FAPI-34 injection were also obtained for dosimetry. Biodistribution of $^{99m}$Tc-FAPI-34 was performed using a gamma counter (Perkin-Elmer), calibrated with known $^{99m}$Tc stock standards. Data was decay-corrected to enable cross-comparison across samples. The biodistribution of caged-MMAE ($^{Cy5}$MSA-pATFB-MMAE) was calculated by quantifying fluorescent imaging taken on the Azure Sapphire FL imaging system using stock standards. Image quantification was performed using ImageJ after subtraction of signal from matched tissues obtained from mice that did not receive $^{Cy5}$MSA-pATFB-MMAE injection to account for background tissue autofluorescence, as previously described(*1,4*). Tissues were then cryopreserved until extraction of MMAE from tissue to estimate drug release was performed.

*RM1.PSMA Prostate Cancer Model*: 5-12 week-old male C57BL/6J mice were inoculated with $2.5 \times 10^5$ RM1.PSMA cells subcutaneously. Roughly 7 days later, once tumors reached approximately ~100mm$^3$ in volume, Alb-caged-exatecan was injected intraperitoneally (5mg/kg free exatecan equivalent, n=2-3 per treatment condition), with $^{177}$Lu-PSMA-617 (74MBq) injected 48h later. After an additional 24 h, tissues were harvested to assess for biodistribution and drug release of Alb-caged-exatecan as outlined below.

***Preclinical dosimetry.***

$^{99m}$Tc-FAPI-34 uptake across tissues over 24h (Fig. 5, Fig. S10B) was fitted with a triexponential decay curve using Python (curve_fit package on SciPy) to estimate the time-integrated activity curves. The absorbed dose was estimated using the MIRD-formalism, with literature-derived S-values(*9,10*).

***Anti-tumor efficacy assay.***

5×10$^5$ TBP3743 cells in PBS were implanted subcutaneously in 5-12 weeks old female B6129SF1/J mice. When tumors ~4-5mm in diameter, mice were randomized and injected intraperitoneally with vehicle control or caged-MMAE (Alb-caged-MMAE, 5 mg/kg equivalent free MMAE) in 100µL PBS. 48h later, 18.5Mbq $^{99m}$Tc-FAPI-34 or $^{177}$Lu-FAPI-04 were injected by tail vein. Subsequent doses of caged-MMAE and $^{99m}$Tc-FAPI-34 were given as indicated in Fig. 6, including 4h after $^{99m}$Tc-FAPI-34 (n = 4-5 mouse per treatment condition, 2 tumors per mouse).

Body weights were monitored, and tumor volumes were calculated using digital caliper measurements and the equation V = length x width$^2$ /2. Prespecified euthanasia criteria included ulceration, tumor size limits, body condition score ≤ 2, or weight loss greater than 20%. Longitudinal tumor growth was plotted as a means for each group until any animals in the group reached the predefined humane experimental endpoint. Toxicity was assayed with complete blood counts, blood chemistry, and histological analysis of the liver and kidneys at the experimental endpoint, performed by the MGH Center for Comparative Medicine Veterinary Pathology Core.

***Tissue extraction of activated prodrug:***

Tissues were harvested at defined time points, weighed, then stored in liquid nitrogen until extraction analysis. Extraction was performed by placing tissue in Lysis buffer II (ThermoFisher, 200µL; with 1X Halt Protease Inhibitor Cocktail). The tissues were finely minced and incubated for 30 minutes on ice, then diluted 4-fold with acetonitrile and centrifuged (5,000 rcf for 5min). The supernatant was filtered through 3kDa MWCO spin filters to remove any remaining tissue fragments and proteins. The flowthrough was analyzed by LCMS or HPLC as above to determine the amount of drug released. Measurements were collected in triplicate and compared to a drug calibration curve to determine concentrations normalized to the mass of the collected tissues (extraction normalization).

***In silico modeling of isotope-dependent drug release efficacy***

Simulations of Alb-caged-MMAE interactions with radionuclides were performed with the Monte Carlo toolkit OpenTOPAS and its extension TOPAS-nBio(*2*) for nanoscopic and microscopic simulations. Simulations were performed using geometries at two spatial scales. (a) First, the vial in which radionuclides and Alb-caged-MMAE were incubated was modeled as a liquid water cylinder of 12.74mm in height and 10mm in diameter, similar to that used for the in vitro experiments described above. Different concentrations of Alb-caged-MMAE were added by randomly placing the corresponding number of macromolecules, considered to approximate spheres of 10 nm in diameter. In turn, radionuclides were assumed to be uniformly distributed in the water so that positions for the emission were randomly selected within the volume of the vial. For each case, $10^7$ histories were simulated, scaling by the actual number of decays to provide each history with an appropriate statistical weight. Using the *g4-livermore* physics model for electromagnetic processes, we calculated the absorbed dose delivered by each radionuclide to the entire vial, considering all the energy-delivering particles and discriminating by each particle. Particles considered were: α-particles, all electrons, electrons producing ionization; electrons undergoing multiple scattering; photons; low-energy electrons (LEEs) defined as those with energy lower than 30 eV; and positrons. Additionally, *virtual* spheres of 10 µm in diameter surrounding the macromolecules were used to store the phase space reaching their surfaces, i.e., the number, type, energy, position, and momentum of all the particles impinging the spheres. This was stored as phase space files and used as the source for (b) microscopic simulations. In this case, a single nanoparticle of 10 nm diameter was set in the middle of a virtual sphere of 10 µm in diameter made of liquid water. The particles recorded at all the bigger spheres from the macroscopic simulations were simulated using the track-structure *geant4-dna (option 2)* physics models, which provide more detailed ionization clouds than the *g4-livermore* physics. We determined the spectrum of electrons reaching macromolecules using the detailed track structure simulations.

#### *Feasibility assessment of radionuclide-mediated drug release in patients*

Dosimetry of cancer lesions in patients was estimated using MIRDcalc(*9*), based on time activity curves inferred from publicly available imaging datasets or published reports of commonly used radiopharmaceutical agents (*11-13*). If required, curves were fitted with a triexponential decay curve to enable estimation of the time-integrated activity curve using Python. Dose per unit activity (Gy/MBq) metrics were calculated for major organs and for tumors of varying sizes and soft tissue composition. Alb-caged-MMAE drug release in tumors was then estimated by applying the expected tumor dose delivered with clinically relevant radionuclide activities to drug release efficiency factors measured in Fig. 2B.

#### *Statistical analysis*

Data were analyzed using GraphPad Prism, MATLAB, and Excel. Normality was tested with the Shapiro-Wilk Test. Specific details of the tests applied are reported in the results. $P<0.05$ was deemed statistically significant.

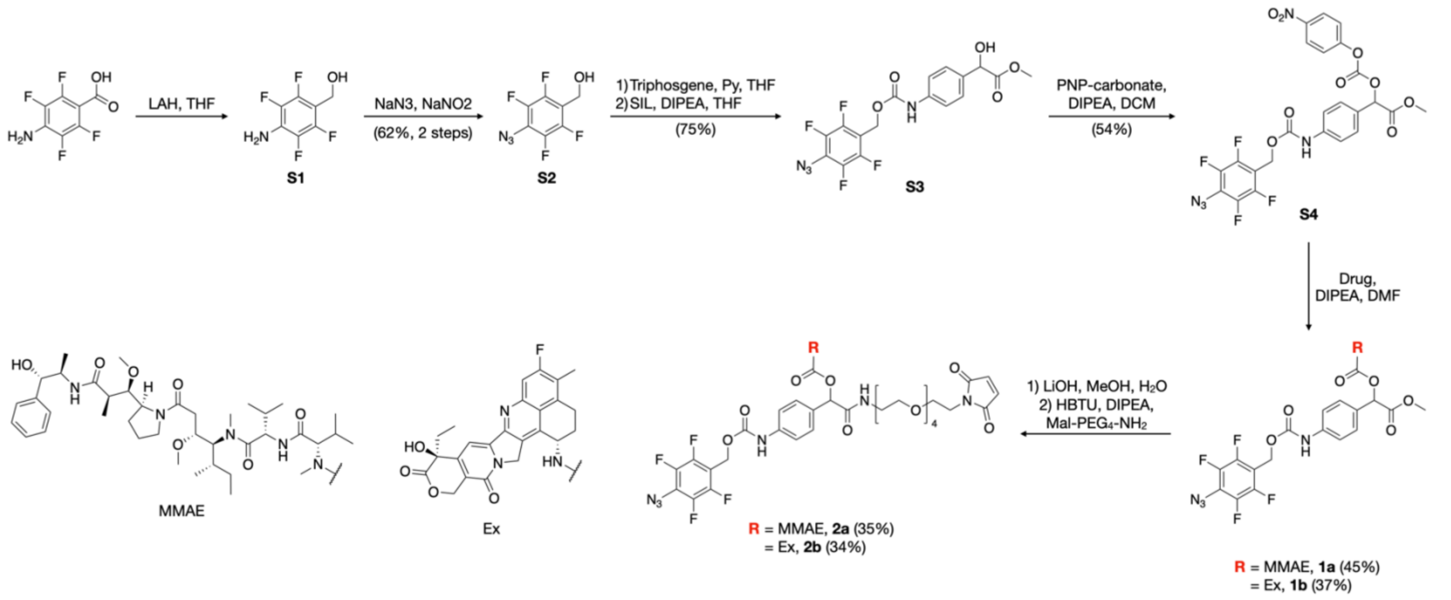


**Scheme S1.** Prodrug synthesis. Compounds **S1-1a** and **2a** were synthesized according to prior literature. (*1,14*)

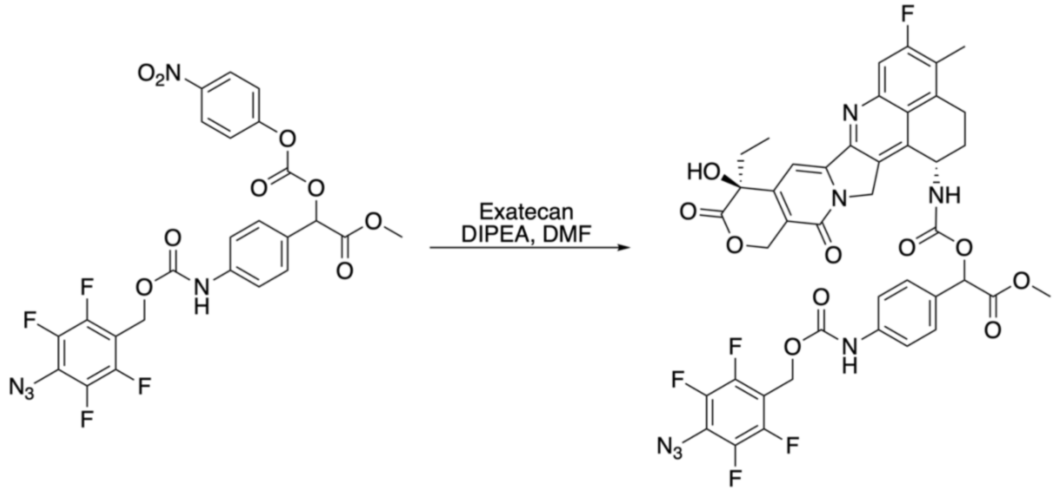

**Scheme S2. Synthesis of pATFB-SIL-exatecan (1b)**. Exatecan (40 mg, 101.9 μmol) was added to a solution of pATFB-SIL-PNP (**S4**, 53.4 mg, 90.0 μmol) in dry DMF (2 mL) and DIPEA (25 μL, 143.5 μmol) was added. This mixture was shaken at room temperature for 18h, then loaded directly onto a reverse-phase column (C18) and purified using a gradient of 5-95% acetonitrile in water (0.1% formic acid). Fractions containing the product were combined and evaporated to provide a brown solid (36.8 mg, 45% yield).

**$^1$H NMR** (400 MHz, $CDCl_3$): δ 8.23 (d, *J* = 9.2 Hz, 1H), 7.63-7.55 (m, 2H), 7.41 (d, *J* = 9.1 Hz, 2H), 7.18 (d, *J* = 8.8 Hz, 1H), 7.09 (d, *J* = 8.2 Hz, 1H), 7.03 (d, *J* = 9.3 Hz, 1H), 6.89 (d, *J* = 9.2 Hz, 1H), 5.90 (s, 1H), 5.62 (dd, *J* = 26.0, 14.4 Hz, 1H), 5.32 (s, 1H), 5.28-5.20 (m, 4H), 5.11 (d, *J* = 16.1 Hz, 1H), 3.77 (s, 1H), 3.75-3.70 (m, 1H), 3.67 (s, 3H), 3.30-3.07 (m, 2H), 2.00 (s, 3H), 1.89-1.77 (m, 2H), 1.38 (d, *J* = 6.7 Hz, 1H), 1.02-0.93 (m, 3H).

**$^{13}$C NMR** (101 MHz, $CDCl_3$): δ 171.7, 168.0, 166.5, 161.7, 160.8, 159.2, 155.6, 153.7, 153.4, 153.0, 150.4, 149.9, 148.6, 148.1, 144.9, 144.2, 143.6, 142.4, 140.3, 139.6, 137.3, 136.3, 133.3, 132.8, 127.7, 126.8, 126.4, 124.1, 123.4, 123.3, 119.8, 119.4, 116.9, 116.4, 116.0, 114.5, 113.8, 112.8, 108.4, 107.9, 107.2, 96.1, 72.7, 70.9, 64.1, 56.1, 52.2, 51.5, 50.7, 48.0, 45.6, 38.9, 34.7, 29.6, 21.6, 18.8, 9.4, 5.9.

**$^{19}$F NMR** (376 MHz, $CDCl_3$): δ -109.6, -142.2, -151.1.

**MS**: m/z calculated for $C_{42}H_{33}F_5N_7O_{10}^+$ $(M+H)^+$ 890.22, found 890.42.

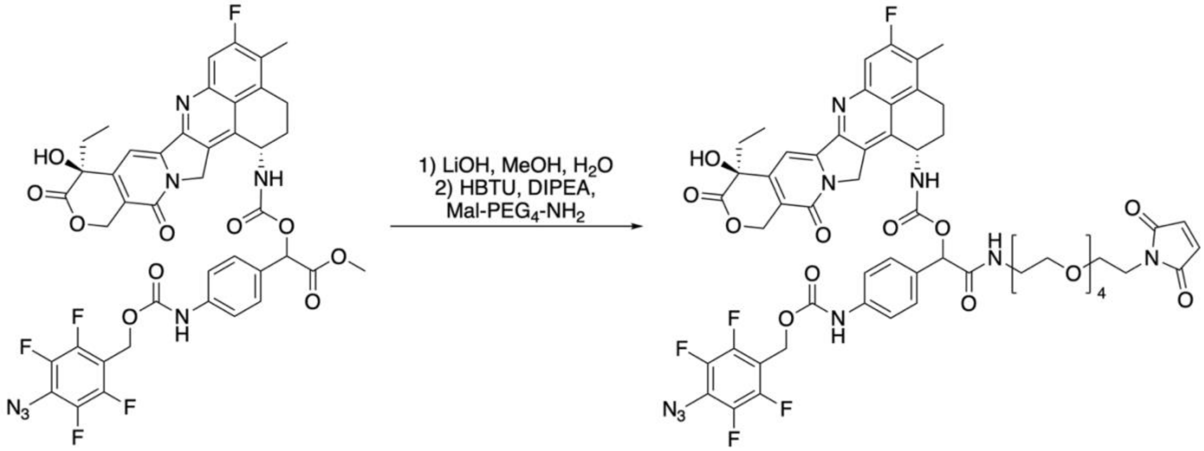


**Scheme S3. Synthesis of pATFB-SIL-Mal-exatecan (2b)**. pATFB-SIL-exatecan (36.8 mg, 24.8 umol) was dissolved in methanol (8 mL) and 0.5M LiOH (2 mL) was added. This mixture was stirred at room temperature for 30 min, then quenched with acidic resin (~1 g). This suspension was stirred for 1 minute, then filtered and washed with methanol. The solvent was removed via rotary evaporation, and the resulting residue was dissolved in DMF (1 mL). HBTU (25 mg, 65.9 umol), Mal-PEG4-amine (100 uL of a 100 mg/mL solution in THF, 31.6 umol), and DIPEA (20 uL, 114.8 umol) were added, and the mixture was stirred for 16h. The reaction mixture was loaded directly onto a reverse-phase column and purified using a gradient of 5-95% acetonitrile in water (0.1% formic acid) to provide the product as a brown solid (22.1 mg, 46% yield).

**$^1$H NMR** (400 MHz, DMSO-$d_6$): δ 8.26 (d, $J$ = 8.3 Hz, 1H), 8.01 (d, $J$ = 8.3 Hz, 2H), 7.95 (s, 1H), 7.40-7.26 (m, 5H), 7.07 (dd, $J$ = 8.4, 4.3 Hz, 1H), 5.30-5.18 (m, 3H), 4.41 (s, 1H), 3.06-2.99 (m, 2H), 2.92-2.77 (m, 12H), 2.73-2.67 (m, 6H), 2.39 (s, 3H), 1.89-1.78 (m, 4H), 1.17-1.13 (m, 5H), 0.85 (s, 3H).

**$^{19}$F NMR** (376 MHz, DMSO-$d_6$): δ -69.2, -71.1, -73.42.

**MS**: m/z calculated for $C_{55}H_{53}F_5N_9O_{15}^+$ $(M+H)^+$ 1174.36, found 1174.53.

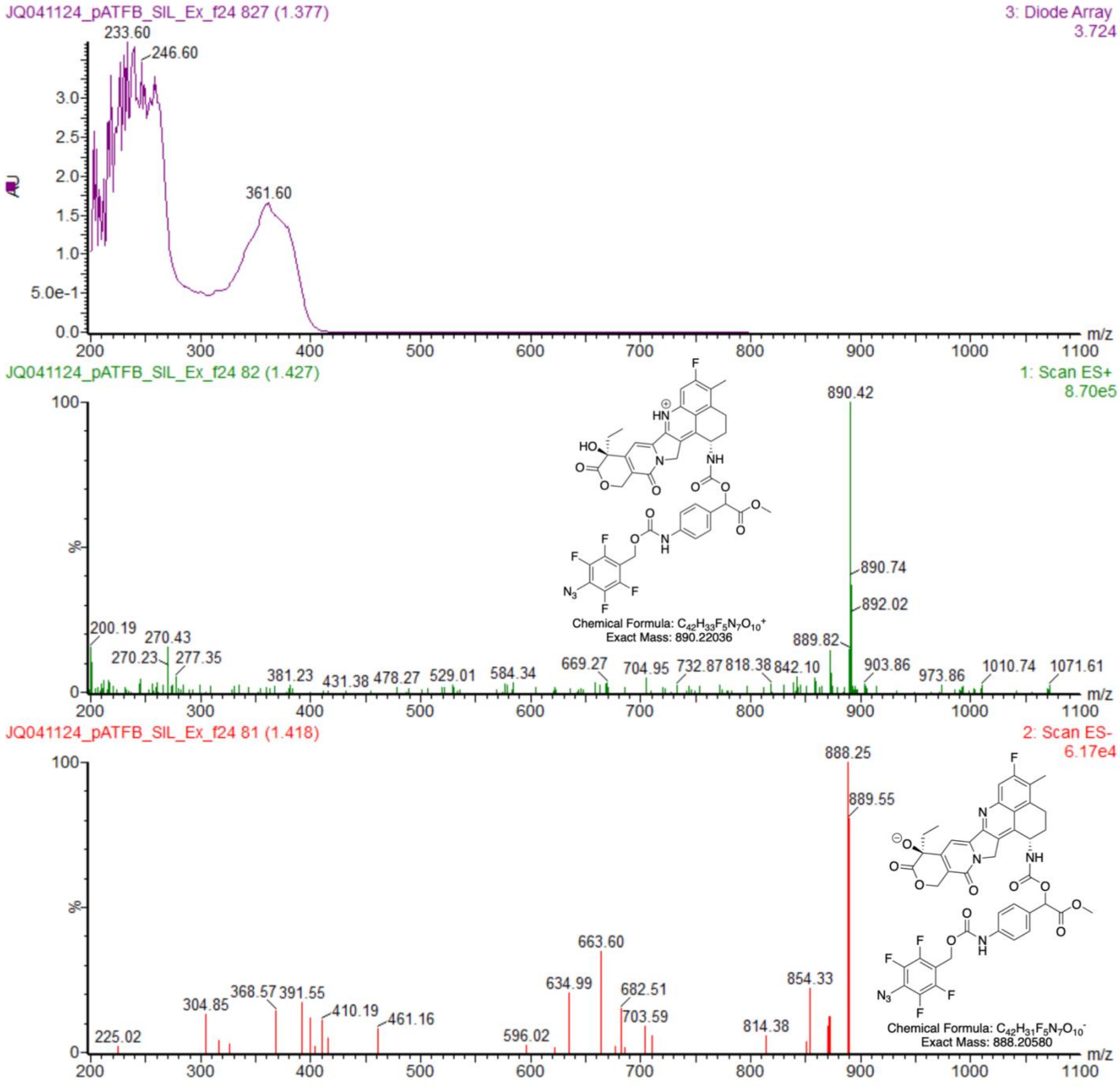


**FIGURE S1.** LCMS spectra of pATFB-SIL-exatecan.

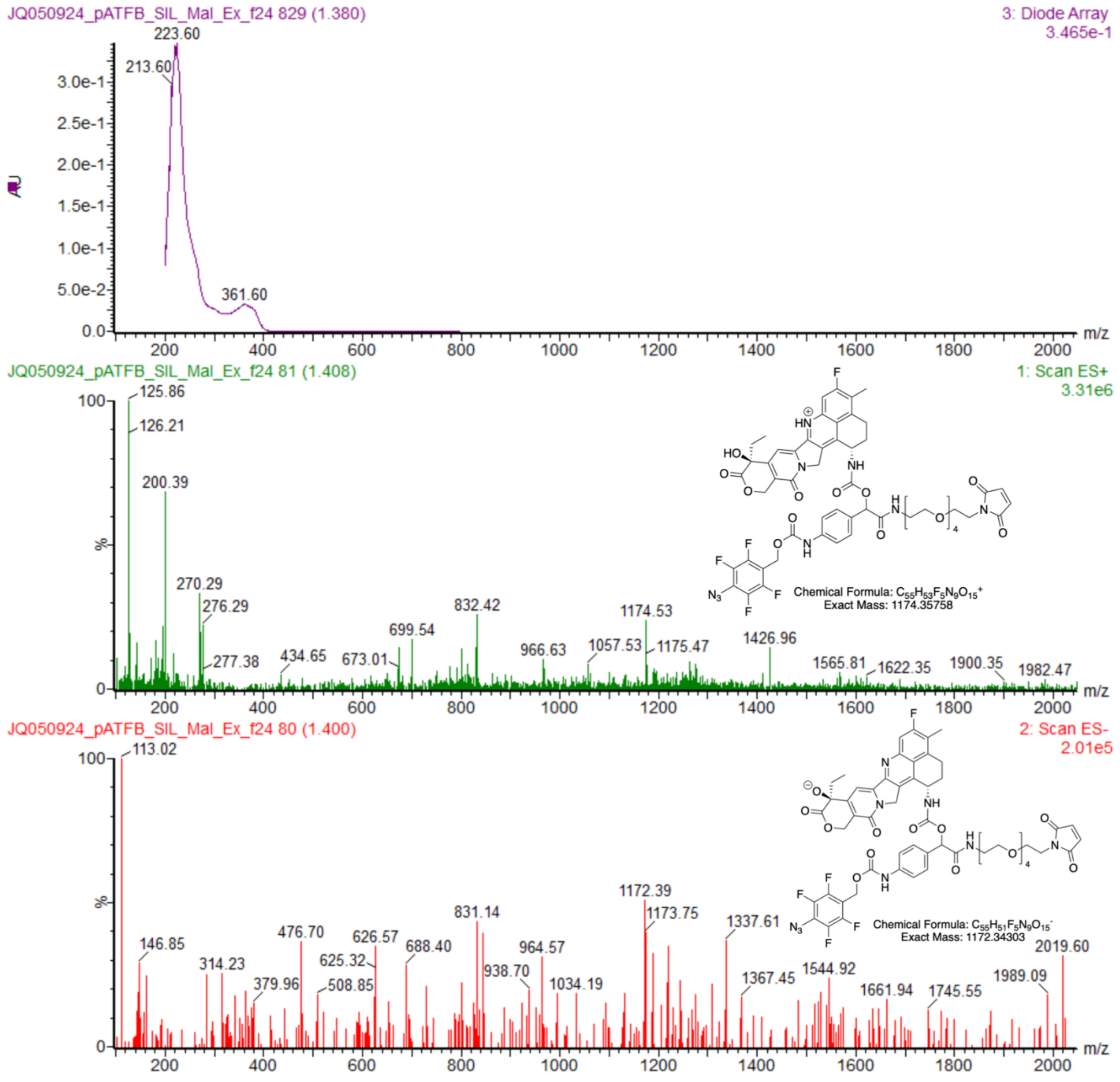


**FIGURE S2**. LCMS spectra of pATFB-SIL-Mal-exatecan.

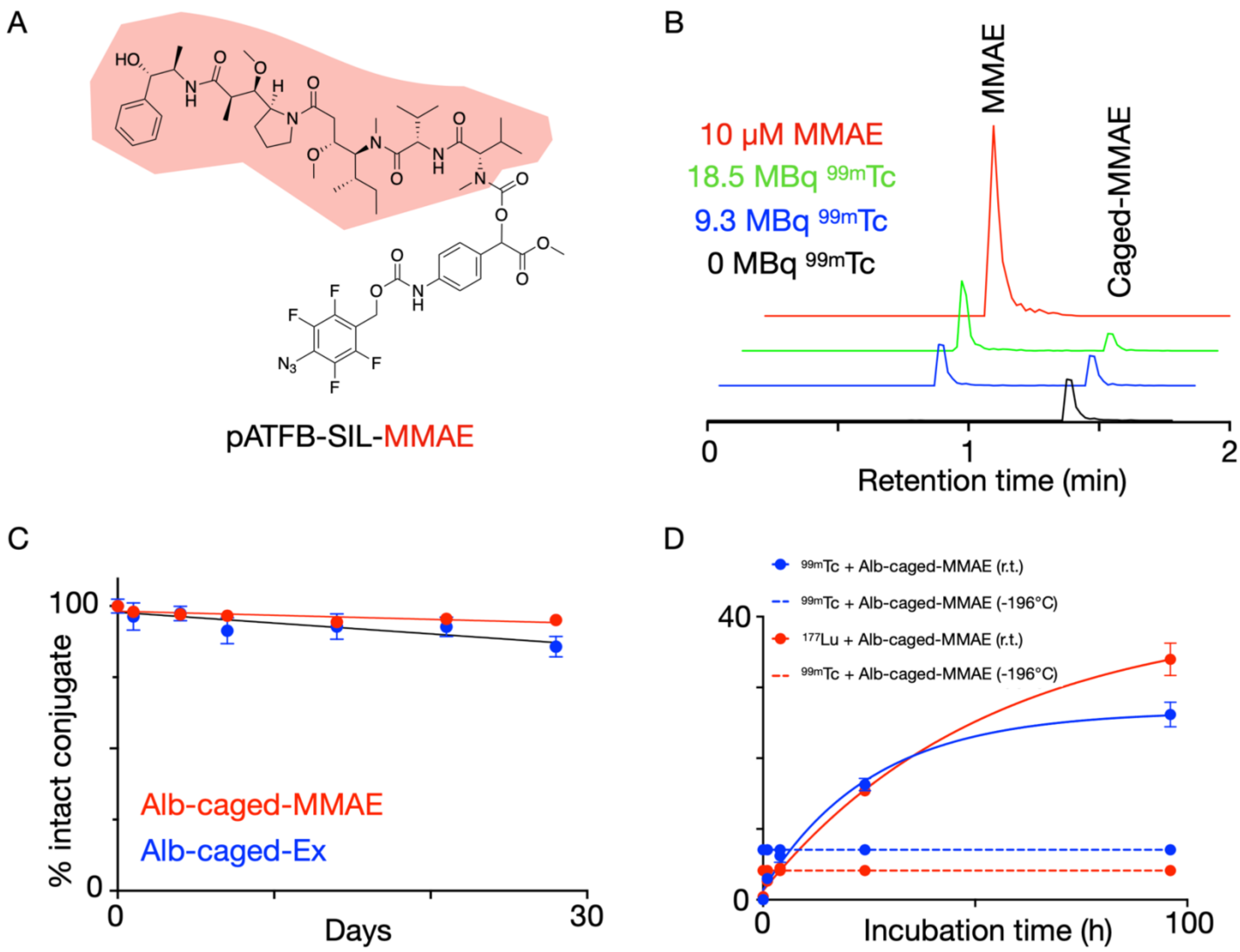


**FIGURE S3.** Caged prodrug stability and release. (A) Caged-MMAE structure: para-azido-2,3,5,6-tetrafluorobenzyl self-immolative linker (pATFB-SIL) conjugated to monomethyl auristatin E (MMAE, red). (B) HPLC trace of caged-MMAE exposed to varying amounts of 99mTc. 10μM MMAE is provided as a comparison (red) (C) Stability of Caged-MMAE and Caged-Exatecan at 4°C in PBS (pH 7.4) over 30 days. (D) Comparison of 99mTc and 177Lu mediated MMAE release from caged-MMAE at room temperature and in liquid nitrogen (-196°C) over 96 h (n=3, mean±S.E.).

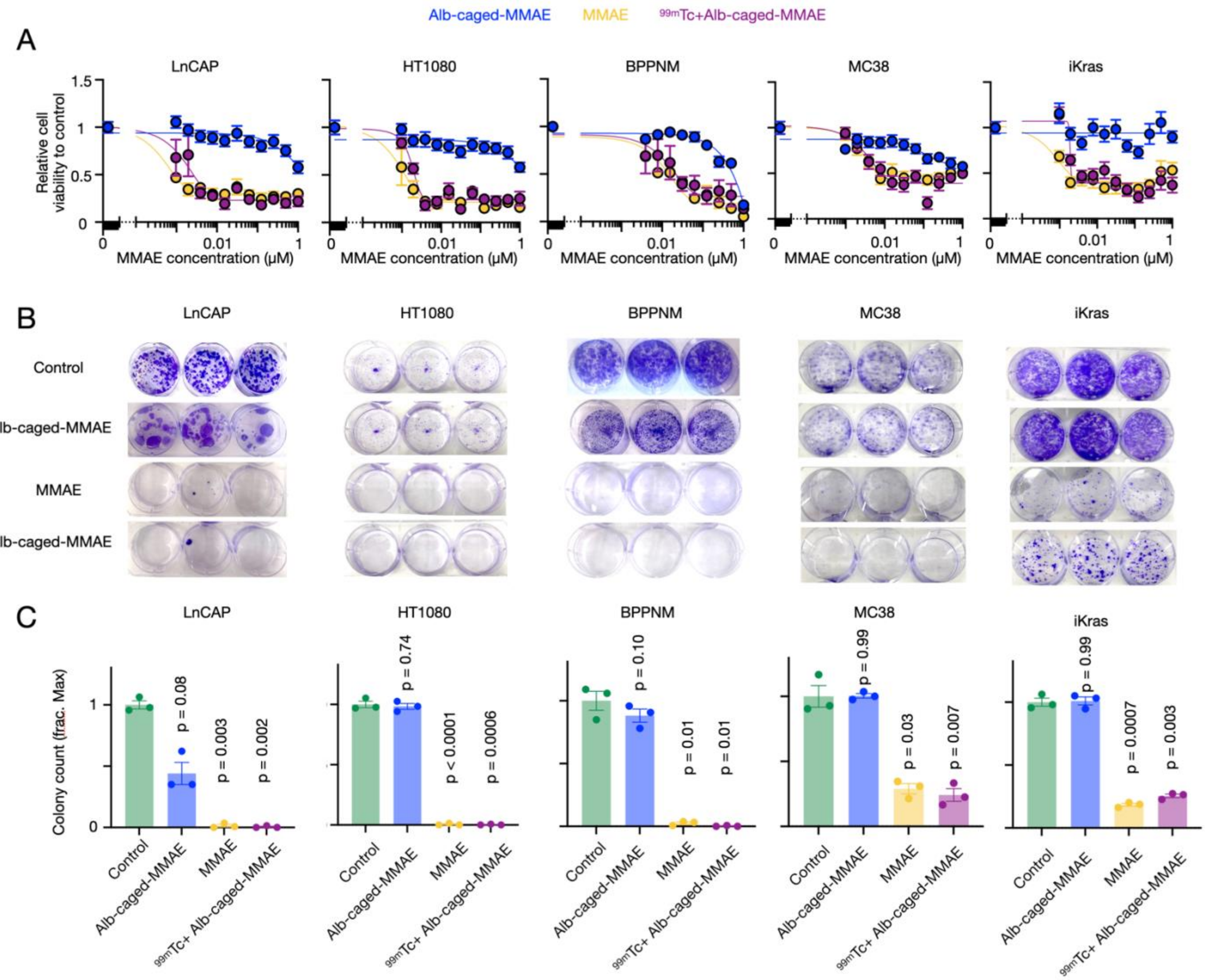


**FIGURE S4.** In vitro assessment of radionuclide-mediated caged-MMAE activation across multiple cell lines. (A) Cytotoxicity assay comparing Alb-caged-MMAE (with and without exposure to $^{99m}$Tc) and free MMAE treatments upon different cancer cell lines, measured in a 72h resazurin-based cytotoxicity assay. Viability compared to untreated control (n=3, mean±S.E.). (B) Representative images and (C) quantification of cancer cell line colony formation (mean±S.E., 2way-ANOVA with Geisser-Greenhouse correction, n=3 per condition).

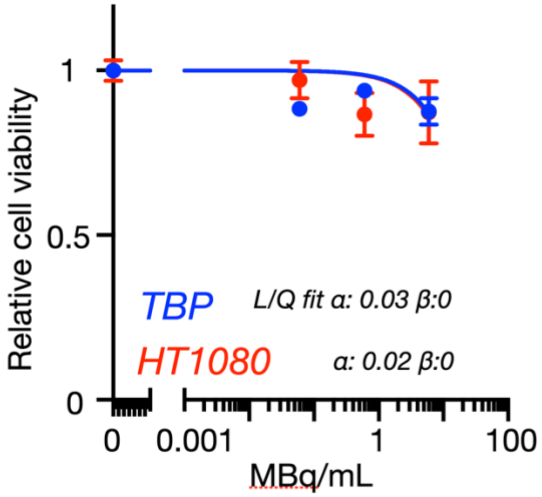


**FIGURE S5**. $^{99m}$Tc cytotoxicity assay. TBP3743 and HT1080 cells were exposed to varying levels of $^{99m}$Tc activities (11.1 MBq/mL, 1.1 MBq/mL and 0.11 MBq/mL) for 72h, followed by viability assessment (n=3, mean±S.E). Curves fitted to the linear quadratic model.

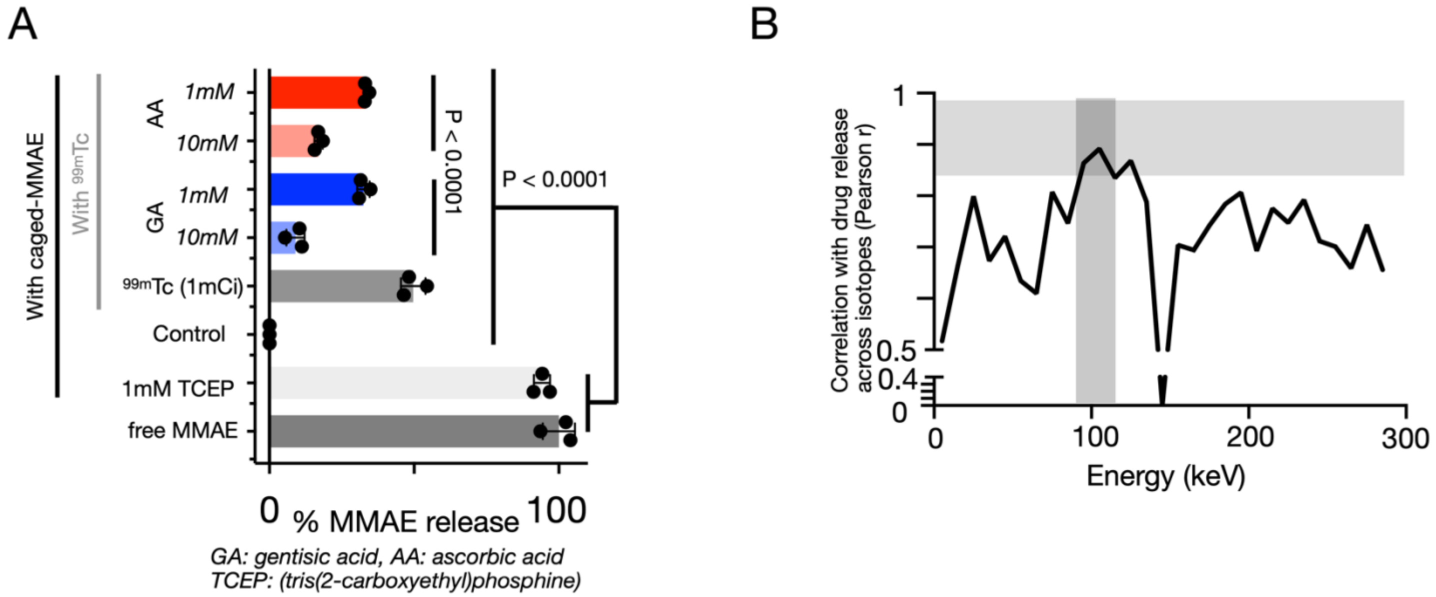


**FIGURE S6**. Identifying mechanisms for RAiDER. (A) The effect of known free radical quenchers gentisic acid (GA) and ascorbic acid (AA) on 99mTc-mediated drug release from caged-MMAE at different concentrations. Release mediated by TCEP, tris(2-carboxyethyl)phosphine, served as a positive control (n=3, mean±S.E., Kruskal-Wallis test). (B) Comparison of drug release and electron generated across different energy windows (0-300keV, 10keV windows) for different radionuclides, measured by Pearson correlation r. The energy window with the highest correlation between electrons generated and drug released (~100keV) was further examined in Fig. 4B (shaded).

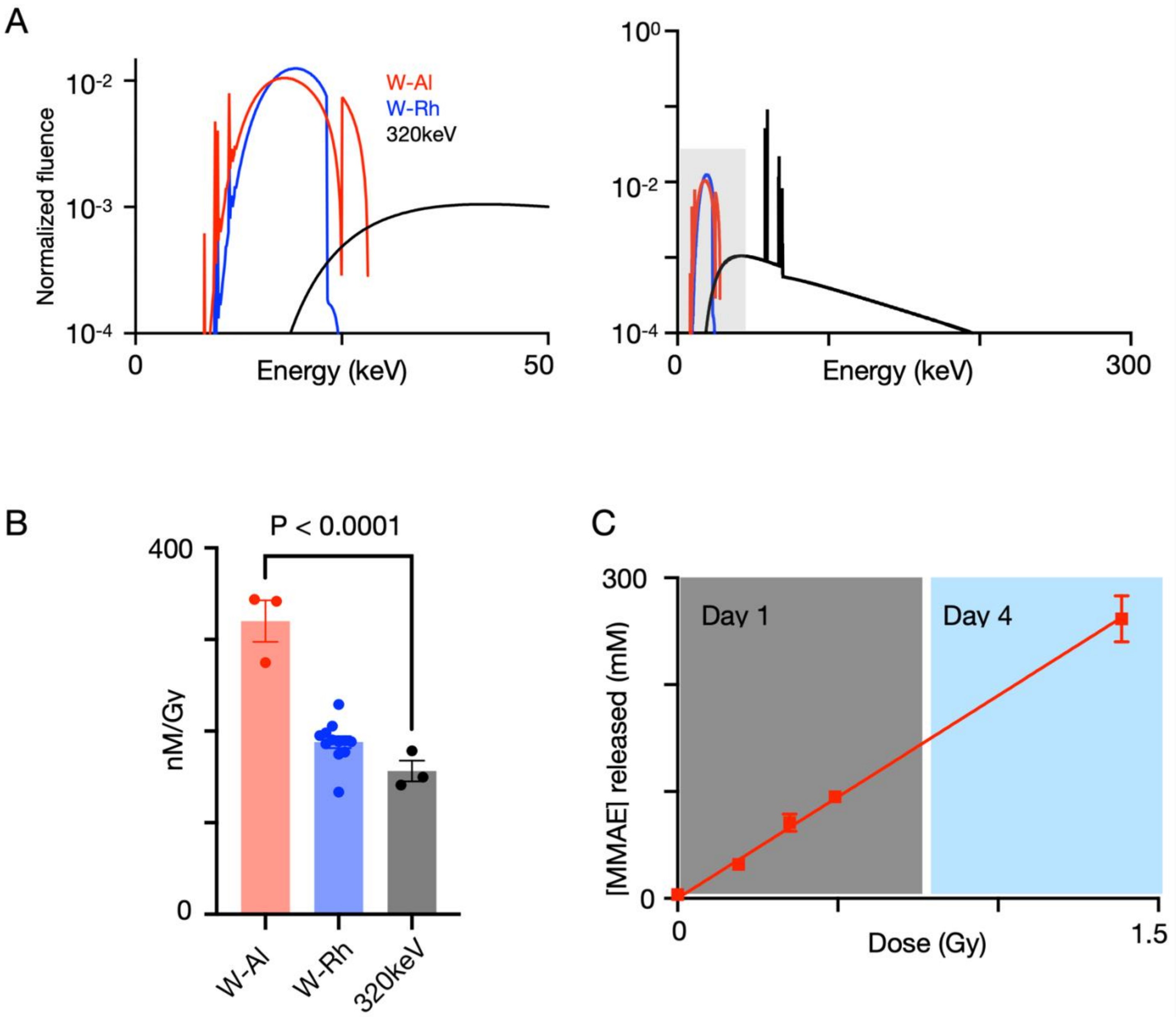


**FIGURE S7**. (A) Mammographic energy spectra using W-Al and W-Rh anode/filter combinations generated by SpekPy(15) show an increase in low-energy photons for W-Al. SpekPy-derived spectra for 320keV X-rays(16) compared to mammography up to 50keV (left) and across 0-300keV (right). Specific SpekPy parameters are noted in Table S3. (B) Experimental drug release efficiency for the different anode/filter combinations shows better W-Al efficiency. This is in keeping with this filter set having more photons emitted in the “low energy” range. Kruskall-Wallis test, n=3. (C) Linearity of mammography-mediated caged-MMAE drug release as a function of dose delivered. Note that high-dose samples (blue) were irradiated over two days, showing linear dose-to-drug release response with this irradiation system (n=3, mean±S.E.).

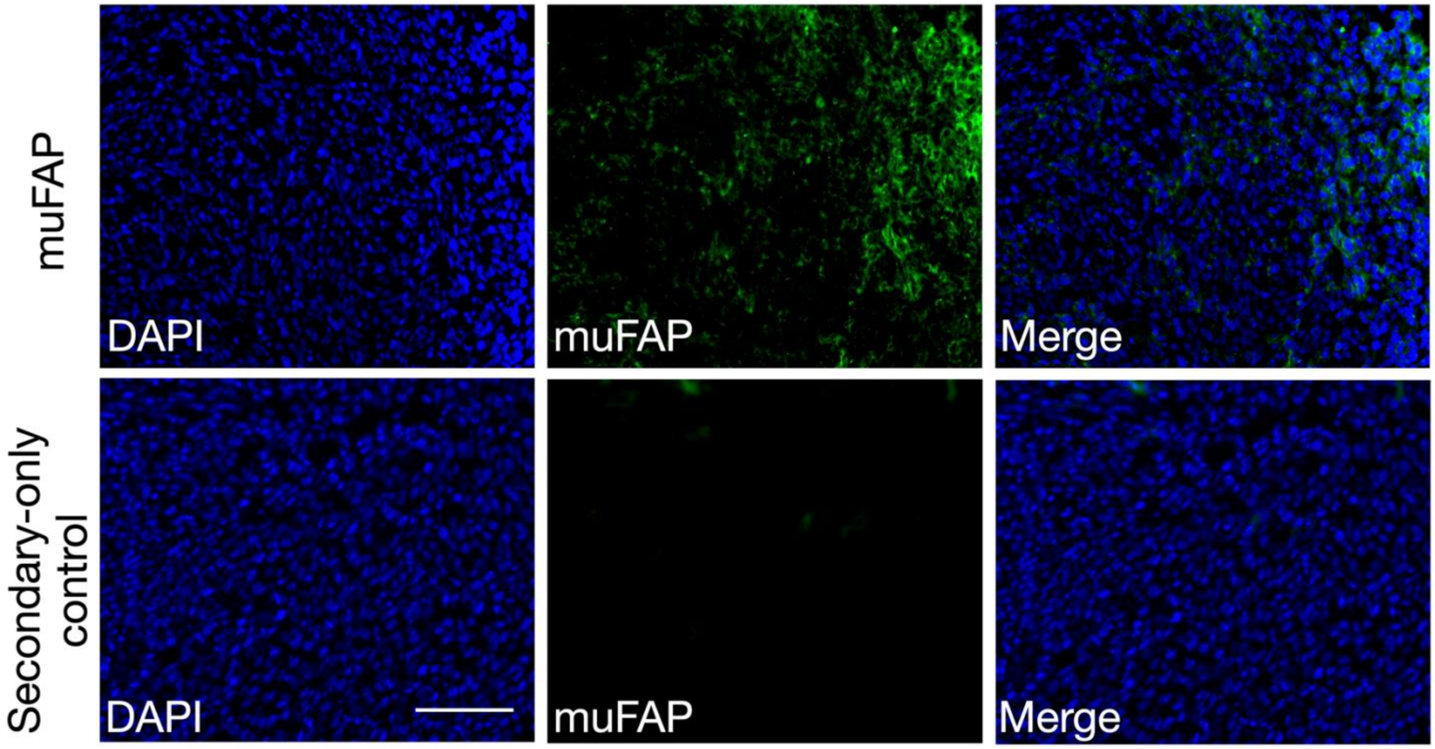


**FIGURE S8.** Murine fibroblast activation protein expression (muFAP, green) in murine TBP3743 tumors. Scale bar = 100μm.

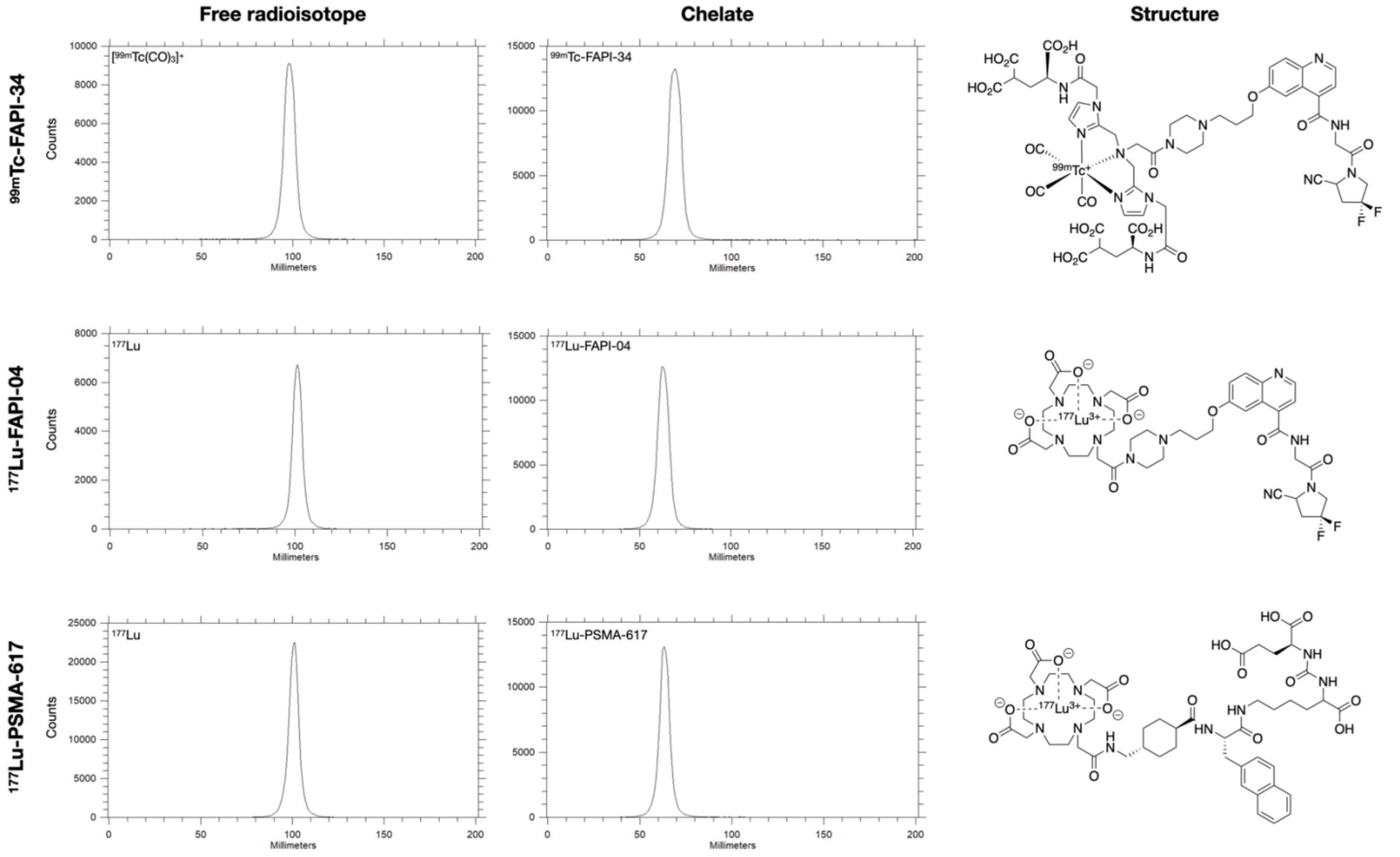


**FIGURE S9**. radioTLC traces for $^{99m}$Tc-FAPI-34, $^{177}$Lu-FAPI-04, and $^{177}$Lu-PSMA-617

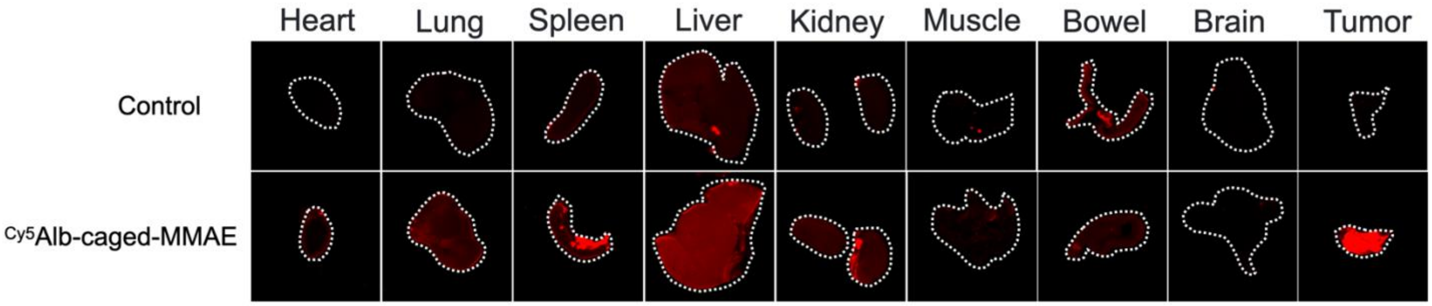


**FIGURE S10.** Representative images showing tissue biodistribution of Cy5Alb-caged-MMAE 48h after intraperitoneal injection (bottom) compared to negative control (top) in B6129SF1/J mice bearing syngeneic TBP3743 tumors and quantified in Fig.5. Tissue borders outlined in white dotted lines (n=3-4 per cohort).

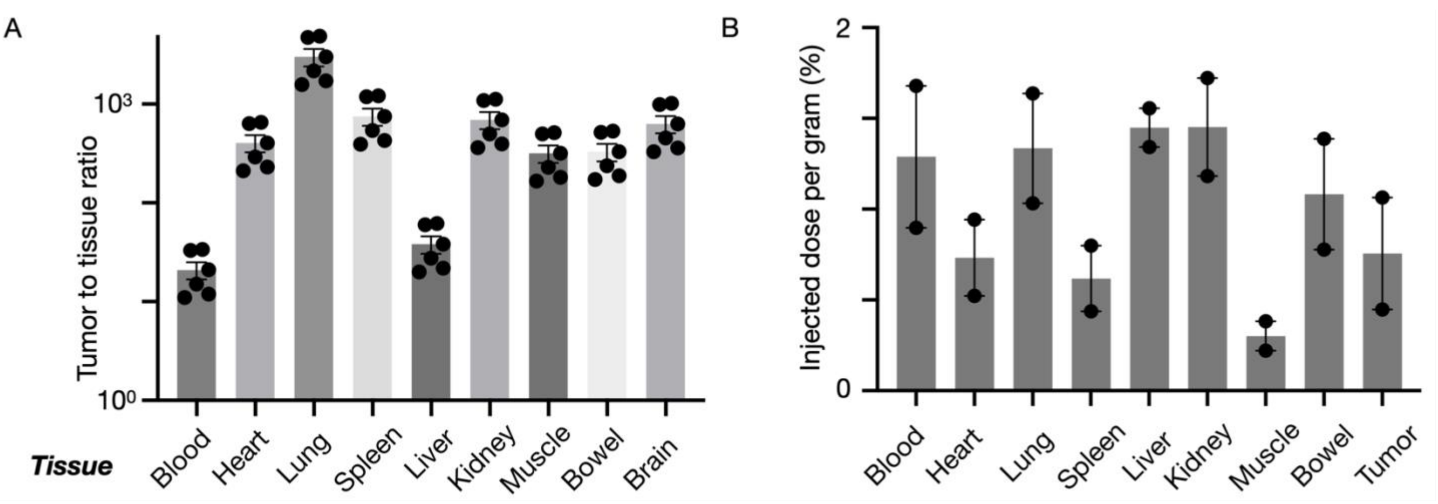


**FIGURE S11.** (A) Tumor-to-tissue ratio of active MMAE in tissues treated with 99mTc-FAPI-34, as depicted in Fig. 5. (B) Biodistribution of 99mTc-FAPI-34 (18.5MBq) in TBP3743 tumor-bearing mice obtained 8 h post-injection (n=2, mean±S.E).

A

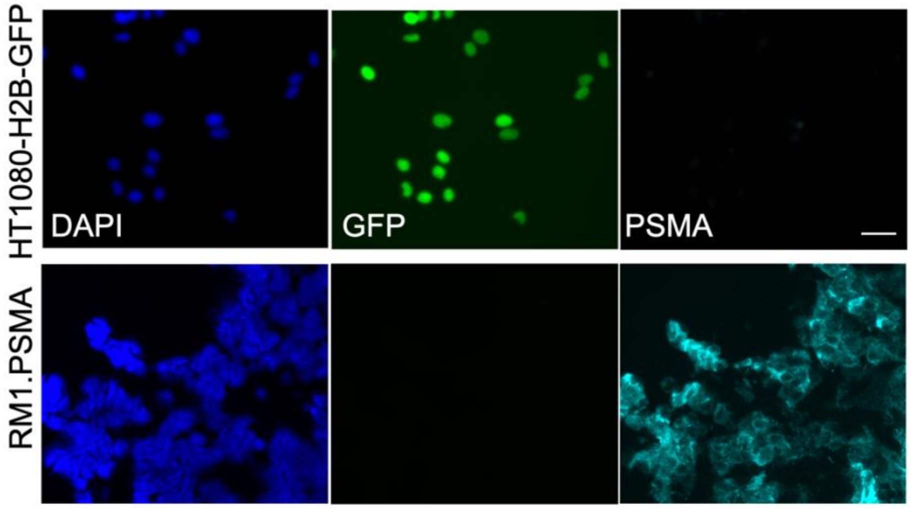


B

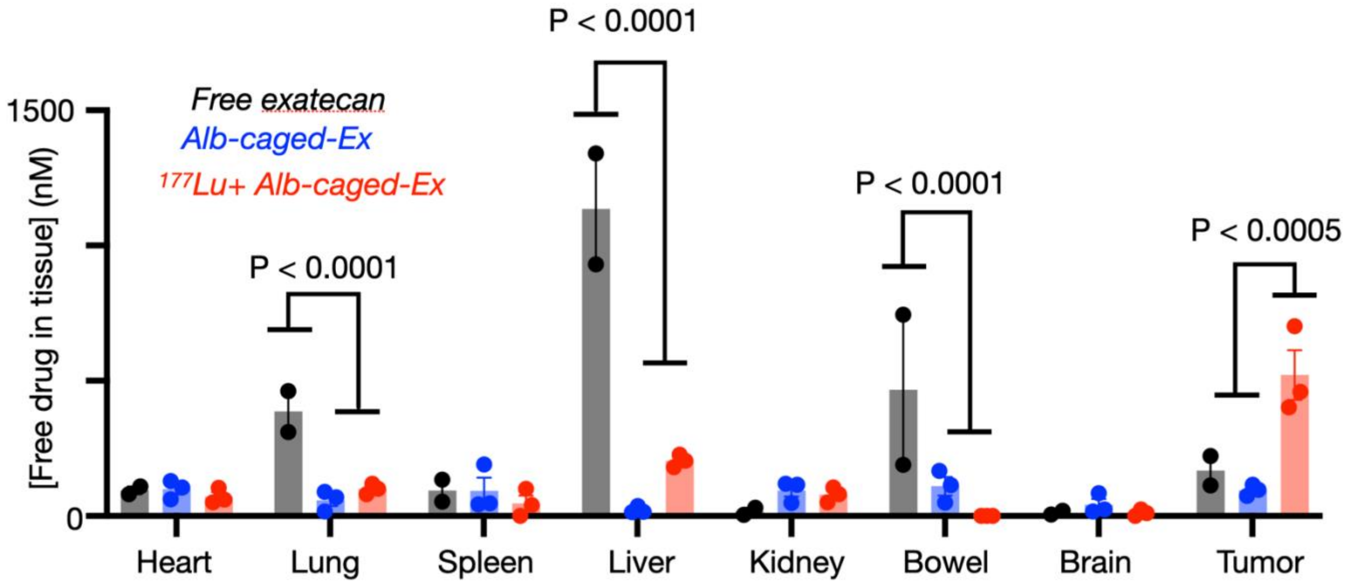


**FIGURE S12.** Drug release mediated by 177Lu-PSMA-617 in prostate cancer. (A) Prostate-specific membrane antigen (PSMA, cyan) expression in RM1.PSMA murine prostate cancer cells, compared to non-PSMA-expressing HT1080-H2B-GFP cells (scale bar = 50μm). (B) RM1.PSMA tumor-bearing mice were injected with Alb-caged-exatecan 48h before 74Mbq 177Lu-PSMA-617. Tissues were harvested 24h later, and exatecan release was assessed. Free exatecan was injected in separate mice for comparison (n=2-3 per treatment, mean±S.E, 2-way-ANOVA with multiple comparisons).

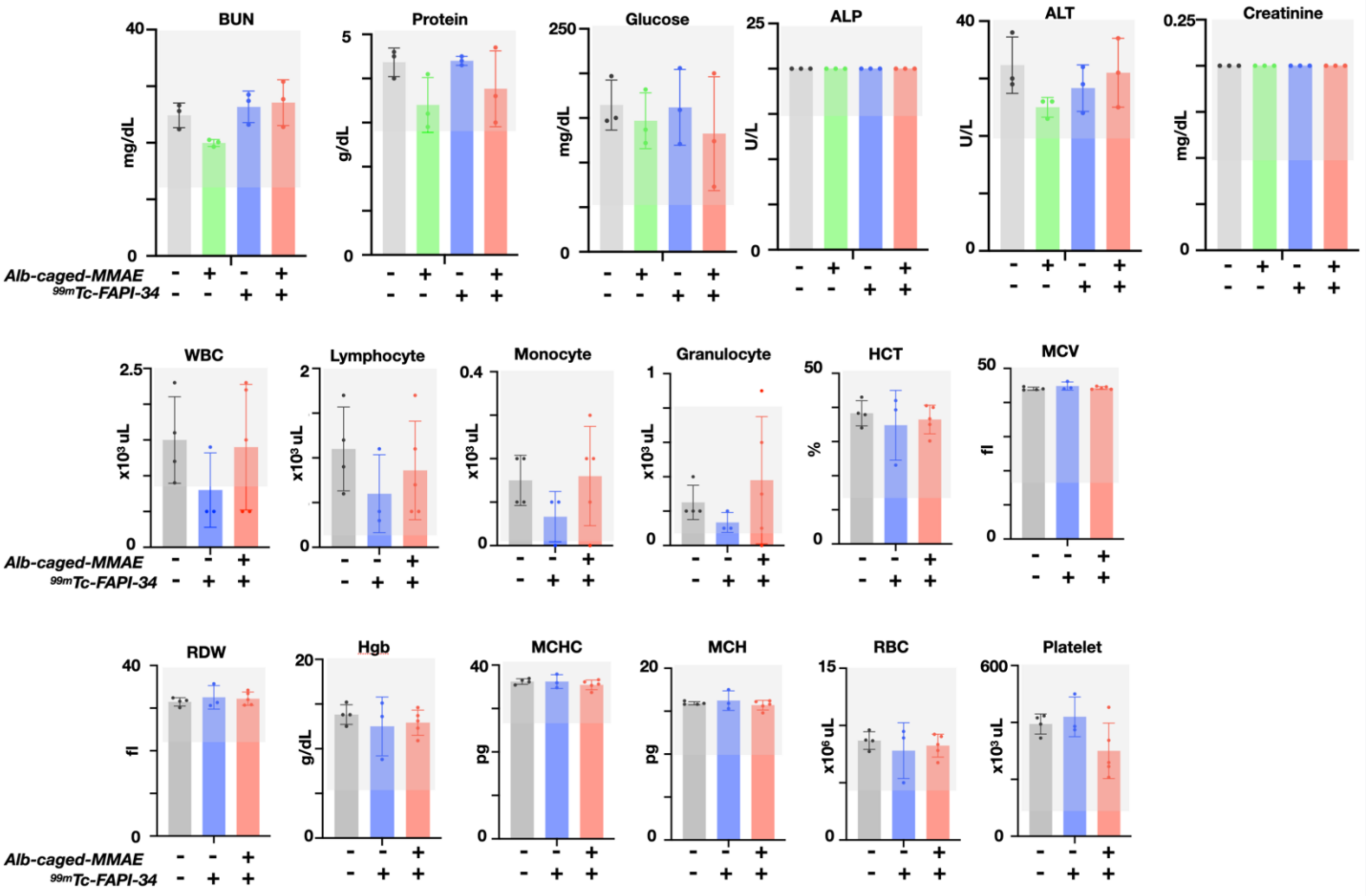


**FIGURE S13**. Blood biomarkers of toxicity during RAiDER. Blood samples were obtained from experimental animals depicted in Fig. 6A at the end of treatment, and a complete blood count (CBC) and complete metabolic panel (CMP) were obtained (mean (S.D)). The shaded box denotes normal ranges. No significant differences were noted across treatment cohorts (Kruskall-Wallis Test, n=3 per treatment).

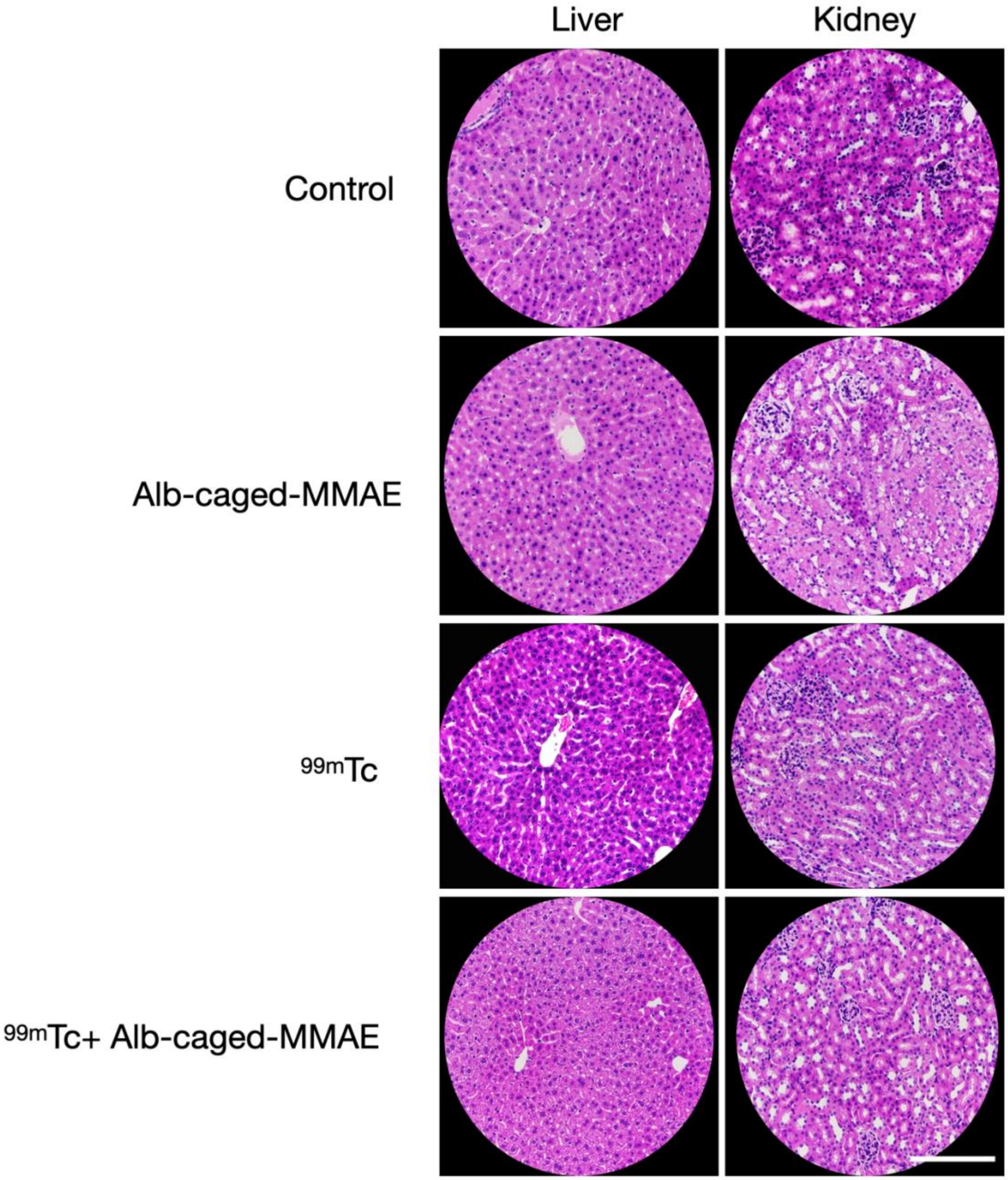


**FIGURE S14.** Representative hematoxylin and eosin staining of liver and kidney tissues across different RAiDER treatments (obtained from subjects depicted in Fig. 6A, 40x). Scale bar 100µm

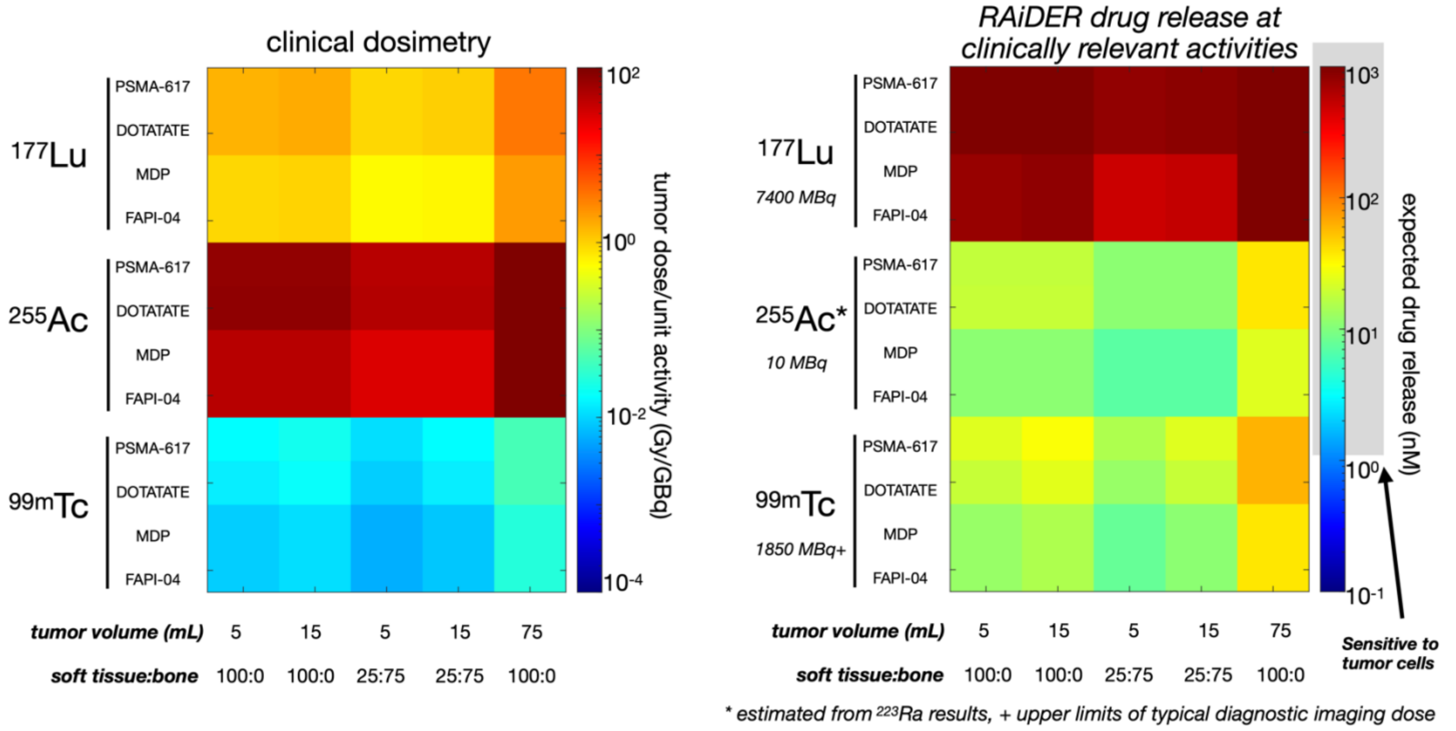


**FIGURE S15.** Computed estimation of RAiDER feasibility in patients. Estimated dosimetry from literature-derived pharmacokinetics and realistic injected doses of commonly used radiopharmaceutical therapies(11-13). MIRCcalc was used for calculations(9). To evaluate whether the tumor absorbed dose would lead to meaningful drug release, the absorbed dose, in Gy/MBq (A), was converted to drug release concentration based on the typical injected dose of the relevant agent in patient and efficiency measures in Fig. 2 (B, Table S5). The assumption is made that enough caged-prodrug can be delivered to the tumor, supported by biodistribution studies(1).

| | Source formulation | Vendor source | Physical $t_{1/2}$ (h) | Decay scheme [ keV/decay, (particles/decay)] | | | | | Major clinical use | Activity tested (Mbq) | Incubation time (h) | Conditions |
|---|---|---|---|---|---|---|---|---|---|---|---|---|
| | | | | Photons | Auger e- | IC e- | Beta (Emax) | Alpha | | | | |
| **$^{18}$F** | [$^{18}$F]$F^-$ | Massachusetts General Hospital Gordon Center positron emission tomography (PET) Core | 1.83 | 511 (1.93) | N/A | N/A | 634 (0.97) | N/A | PET | 3.70E+01<br>8.88E+01<br>1.11E+02 | 24 | room temperature |
| **$^{64}$Cu** | [$^{64}$Cu]$CuCl_2$ | Department of Medical Physics, University of Wisconsin (Madison, WI) | 12.7 | 511 (0.34)<br>7.4 (0.14)<br>(x-rays) | 2.5 (0.8) | N/A | 579 (0.39)<br>653 (0.17) | N/A | PET | 5.55E+00<br>1.11E+01<br>1.85E+01 | 168 | room temperature |
| **$^{68}$Ga** | [$^{68}$Ga]$GaCl_3$ | Galli Eo$^{68}$Ge/$^{68}$Ga generator (IRE ELiT) | 1.1 | 511 (1.78) | 2.7 (0.19) | 9 (0.04) | 1899 (0.89) | N/A | PET | 3.70E+01<br>1.11E+02 | 24 | room temperature |
| **$^{99m}$Tc** | $Na^+$[$^{99m}$Tc]$TcO_4^-$ | $^{99}$Mo/$^{99m}$Tc generator, TECHNELITE®, Lantheus Medical Imaging, Inc. | 6 | 140 (0.89) | 0.9 (4.4) | 15.2 (1.1) | N/A | N/A | SPECT | 9.25E+00<br>1.85E+01<br>3.70E+01 | 1, 4, 24, 96, 120, 168 | room temperature |
| **$^{177}$Lu** | [$^{177}$Lu]$LuCl_2$ | Eckert & Ziegler Medical/SHINE, no carrier added | 160.8 | 208 (0.11)<br>113 (0.06) | 6.2 (0.09) | 83 (0.13) | 497 (0.79)<br>384 (0.09)<br>176 (0.12) | N/A | RPT | 5.55E+00<br>1.11E+01 | 1, 4, 24, 96, 120, 168 | room temperature<br>-196°C |
| **$^{223}$Ra** | [$^{223}$Ra]$RaCl_2$ | Massachusetts General Hospital Radiation Safety Office | 273.6 | 84 (0.41)<br>270 (0.24)<br>350 (0.13)<br>154 (0.09) | 8.7 (0.3) | 3-24 (0.65) | 470 (1.9) | 5000-7500 (0.95) | RPT | 4.63E-03<br>1.85E-02<br>7.40E-02 | 168 | room temperature<br>-196°C |

*SPECT: single photon emission computed tomography, PET: positron emission tomography, RPT: radiopharmaceutical therapy, IC: internal conversion*

**Table S1.** Properties of isotopes and in vitro experimental conditions used in this study.

| ***Cell line*** | ***Tumor Type*** | ***IC50 (nM)*** | | |
|---|---|---|---|---|
| | | ***MMAE*** | ***MSA-pAFTB-MMAE*** | ***$^{99m}$Tc + MSA-pAFTB-MMAE*** |
| HT1080 | *fibrosarcoma* | 2.4 | 2000 | 4.5 |
| TBP | *anaplastic thyroid cancer* | 4.9 | 500 | 2.9 |
| BBPNM | *ovarian cancer* | 29 | 460 | 16 |
| LnCAP | *prostate cancer* | 1.8 | 1200 | 3.4 |
| MC38 | *colorectal cancer* | 30 | 640 | 17 |
| iKRAS | *pancreatic cancer* | 4.2 | 8300 | 13 |

**Table S2.** Estimated 50% proliferation/cytotoxicity inhibition (IC50) values from viability curves generated in Fig. 3A and Fig. S4 across different cancer cell lines across Alb-caged-MMAE, free MMAE, and 99mTc+Alb-caged-MMAE treatments.

| | W-Rh | W-Al | 320 kV* |
|---|---|---|---|
| ***Physics*** | kqp | kqp | kqp |
| ***Attenuation data*** | Penelope | Penelope | Penelope |
| ***Energy bin (keV)*** | 0.1 | 0.1 | 0.1 |
| ***Target material*** | W | W | W |
| ***Anode angle (degrees)*** | 16 | 16 | 30 |
| ***kV*** | 25 | 25 | 300 |
| ***x (cm)*** | 0 | 0 | 0 |
| ***y (cm)*** | 0 | 0 | 0 |
| ***z (cm)*** | 70 | 70 | 70 |
| ***mAs*** | 17875 | 9500 | 2250 |
| ***Bremsstrahlung*** | true | true | true |
| ***Characteristic x-ray*** | true | true | true |
| ***Filter*** | Rh | Al | Al |
| ***Filter thickness (mm)*** | 0.0563 | 0.723 | 2 |

*Matched to XRAD320 (16)

**Table S3.** SpekPy parameters for simulations.

| | $^{99m}$Tc-FAPI-34 | |
|---|---|---|
| | ***Gy/MBq*** | ***Absorbed Dose (Gy)**** |
| **blood** | 2.70E-03 | 5.00E-02 |
| **heart** | 2.70E-03 | 4.99E-02 |
| **lung** | 5.37E-03 | 9.94E-02 |
| **spleen** | 3.09E-03 | 5.72E-02 |
| **liver** | 6.40E-03 | 1.18E-01 |
| **kidney** | 6.78E-03 | 1.25E-01 |
| **muscle** | 1.43E-03 | 2.65E-02 |
| **bowel** | 5.43E-03 | 1.01E-01 |
| **tumor┼** | 6.47E-03 | 1.20E-01 |

**Based on 18.5 MBq injected*

*Expected tumor [MMAE] based on in vitro results: 1.45E02 nM*

**Table S4.** Estimated dosimetry of 99mTc-FAPI-34 based on time-injected activity curves derived from Fig. 5 and Fig. S11 using MIRD formalism(9).

| Pharmaceutical | Isotope | Size (mL) | % Soft tissue | mGy/MBq | Clinical dosing (MBq) | Gy | Estimated drug release (nM)* |
|---|---|---|---|---|---|---|---|
| PSMA-617 | $^{177}Lu$ | 5 | 100 | 1.53 | 7400 | 11.32 | 1308 |
| | | 15 | 100 | 1.66 | 7400 | 12.28 | 1419 |
| | | 6 | 25 | 0.96 | 7400 | 7.11 | 821 |
| | | 15 | 25 | 1.02 | 7400 | 7.55 | 872 |
| | | 75 | 100 | 3.47 | 7400 | 25.69 | 2968 |
| | $^{225}Ac$ | 5 | 100 | 75.10 | 20 | 1.50 | 18 |
| | | 15 | 100 | 75.50 | 20 | 1.51 | 18 |
| | | 6 | 25 | 45.70 | 20 | 0.91 | 11 |
| | | 15 | 25 | 45.80 | 20 | 0.92 | 11 |
| | | 75 | 100 | 157.90 | 10 | 3.16 | 38 |
| | $^{99m}Tc$ | 5 | 100 | 0.02 | 1110 | 0.02 | 23 |
| | | 15 | 100 | 0.02 | 1110 | 0.02 | 30 |
| | | 6 | 25 | 0.01 | 1110 | 0.01 | 15 |
| | | 15 | 25 | 0.02 | 1110 | 0.02 | 23 |
| | | 75 | 100 | 0.05 | 1110 | 0.05 | 62 |
| DOTATATE | $^{177}Lu$ | 5 | 100 | 1.53 | 7400 | 11.32 | 1308 |
| | | 15 | 100 | 1.66 | 7400 | 12.28 | 1419 |
| | | 6 | 25 | 0.96 | 7400 | 7.11 | 821 |
| | | 15 | 25 | 1.02 | 7400 | 7.55 | 872 |
| | | 75 | 100 | 3.47 | 7400 | 25.69 | 2968 |
| | $^{225}Ac$ | 5 | 100 | 77.50 | 20 | 1.55 | 19 |
| | | 15 | 100 | 77.90 | 20 | 1.56 | 19 |
| | | 6 | 25 | 47.20 | 20 | 0.94 | 11 |
| | | 15 | 25 | 47.30 | 20 | 0.95 | 11 |
| | | 75 | 100 | 157.90 | 20 | 3.16 | 38 |
| | $^{99m}Tc$ | 5 | 100 | 0.01 | 1110 | 0.02 | 19 |
| | | 15 | 100 | 0.02 | 1110 | 0.02 | 24 |
| | | 6 | 25 | 0.01 | 1110 | 0.01 | 12 |
| | | 15 | 25 | 0.01 | 1110 | 0.02 | 19 |
| | | 75 | 100 | 0.05 | 1110 | 0.05 | 62 |
| MDP | $^{177}Lu$ | 5 | 100 | 0.92 | 7400 | 6.81 | 786 |
| | | 15 | 100 | 1.00 | 7400 | 7.40 | 855 |
| | | 6 | 25 | 0.58 | 7400 | 4.27 | 493 |
| | | 15 | 25 | 0.62 | 7400 | 4.57 | 527 |
| | | 75 | 100 | 2.09 | 7400 | 15.48 | 1788 |
| | $^{225}Ac$ | 5 | 100 | 46.80 | 20 | 0.94 | 11 |
| | | 15 | 100 | 46.90 | 20 | 0.94 | 11 |
| | | 6 | 25 | 28.40 | 20 | 0.57 | 7 |
| | | 15 | 25 | 28.50 | 20 | 0.57 | 7 |
| | | 75 | 100 | 95.10 | 20 | 1.90 | 23 |
| | $^{99m}Tc$ | 5 | 100 | 0.01 | 1110 | 0.01 | 12 |
| | | 15 | 100 | 0.01 | 1110 | 0.01 | 15 |
| | | 6 | 25 | 0.01 | 1110 | 0.01 | 8 |
| | | 15 | 25 | 0.01 | 1110 | 0.01 | 11 |
| | | 75 | 100 | 0.03 | 1110 | 0.03 | 38 |
| FAPI-04 | $^{177}Lu$ | 5 | 100 | 0.92 | 7400 | 6.81 | 786 |
| | | 15 | 100 | 1.00 | 7400 | 7.40 | 855 |
| | | 6 | 25 | 0.58 | 7400 | 4.27 | 493 |
| | | 15 | 25 | 0.62 | 7400 | 4.57 | 527 |
| | | 75 | 100 | 2.09 | 7400 | 15.48 | 1788 |
| | $^{225}Ac$ | 5 | 100 | 46.80 | 20 | 0.94 | 11 |
| | | 15 | 100 | 46.90 | 20 | 0.94 | 11 |
| | | 6 | 25 | 28.40 | 20 | 0.57 | 7 |
| | | 15 | 25 | 28.50 | 20 | 0.57 | 7 |
| | | 75 | 100 | 95.10 | 20 | 1.90 | 23 |
| | $^{99m}Tc$ | 5 | 100 | 0.01 | 1110 | 0.01 | 12 |
| | | 15 | 100 | 0.01 | 1110 | 0.01 | 15 |
| | | 6 | 25 | 0.01 | 1110 | 0.01 | 8 |
| | | 15 | 25 | 0.01 | 1110 | 0.01 | 11 |
| | | 75 | 100 | 0.03 | 1110 | 0.03 | 38 |

** Estimated from in vitro derived values from Figure 2*

**Table S5.** MIRDCalc(9) estimation of clinically realistic dosimetry and expected drug release across different isotopes/radiopharmaceutical agents as depicted in Fig. S15.
